# Effect of sublethal prenatal endotoxaemia on murine placental transport systems and lipid homeostasis

**DOI:** 10.1101/2020.08.04.236745

**Authors:** MW Reginatto, KN Fontes, VRS Monteiro, NL Silva, CBV Andrade, HR Gomes, GE Imperio, FF Bloise, G Kluck, GC Atella, SG Matthews, E Bloise, TM Ortiga-Carvalho

## Abstract

Infection alters the expression of transporters that mediate the placental exchange of xenobiotics, lipids and cytokines. We hypothesized that lipopolysaccharide (LPS) modifies the expression of placental transport systems and lipid homeostasis. LPS (150 μg/kg; i.p.) treatments were administered for 4 h or 24 h, animals were euthanized at gestational days (GD) 15.5 or 18.5, and maternal blood, foetuses and placentae were collected. Increased rates of foetal demise were observed at GD15.5 following LPS treatment, whereas at GD18.5, high rates of early labour occurred and were associated with distinct proinflammatory responses. LPS did not alter ABC transporter mRNA expression but decreased Fabppm at GD15.5 (LPS-4 h) and increased Fat/Cd36 lipid transporter mRNA at GD18.5 (LPS-4 h). At the protein level, breast cancer-related protein (BCRP) and Abcg1 levels were decreased in the placental labyrinth zone (Lz) at GD15.5, whereas P-glycoprotein (P-gp) and Bcrp Lz-immunostaining was decreased at GD18.5. In the placental junctional zone (Jz), P-gp, Bcrp and Abcg1 levels were higher at GD18.5. Specific maternal plasma and placental changes in triacylglycerol, free fatty acid, cholesterol, cholesterol ester and monoacylglycerol levels were detected in a gestational age-dependent manner. In conclusion, LPS-induced foetal death and early labour were associated with altered placental ABC and lipid transporter expression and deranged maternal plasma and placental lipid homeostasis. These changes likely modify foetal xenobiotic exposure and placental lipid exchange in cases of bacterial infection.

## Introduction

Global estimates indicate that more than 15 million babies are born preterm every year [1]. Preterm birth (PTB) occurs at higher rates in low- and middle-income countries and may range from 5 to 18% of all pregnancies worldwide [2]. Of particular importance, low-income countries have a higher incidence of histological chorioamnionitis (HCA)-related genitourinary (retrograde or ascending bacterial) and malarial (haematogenous) infections. These infections may lead to severe systemic and placental inflammatory response [3–5] and become important triggers of inflammatory PTB pathways [1,6,7]. Other routes/risk factors for infection-associated PTB include maternal periodontal disease, transplacental transfer of pathogens, iatrogenic infection from complicating amniocentesis or chorionic villous sampling [7,8].

The most common microbes observed in HCA are the gram-negative bacteria Ureaplasma urealyticum, Mycoplasma hominis and Escherichia coli [7]. Lipopolysaccharide (LPS), an endotoxin enriched in the cell wall of gram-negative bacteria, binds Toll-like receptor 4 (Tlr-4) and is widely used to model gram-negative bacterial infections [9]. Studies in mice have shown that LPS exposure alters the foetal-placental unit in a gestational age-dependent manner. In early pregnancy, it induces embryonic loss [10] or miscarriage [11], whereas in mid-pregnancy, it causes foetal death, foetal growth restriction [12] and foetal brain injury [13].

Infection and inflammation have the potential to disrupt the syncytiotrophoblast barrier by modulating the expression and function of ATP-binding cassette (ABC) transporters [14–16]. These proteins are active transmembrane efflux transport systems that control the biodistribution of clinically relevant endogenous and exogenous substrates across the maternal-foetal interface. Examples of endogenous substrates include nutrients (cholesterol and other lipids), metabolites (bilirubin- and bile salt-conjugated compounds and oxysterols), steroid hormones (glucocorticoids, mineralocorticoids, oestrogens, progestogens, and androgens) and immunological factors (cytokines and chemokines). Examples of exogenous substrates include therapeutic drugs (antibiotics, antiretrovirals, synthetic glucocorticoids and NSAIDs) and environmental toxins (organochlorine and organophosphorus pesticides, ivermectin and bisphenol A). As a result, ABC transporters limit the transfer of potentially harmful substrates to the foetus and control transplacental passage of nutrients (mostly lipids) and other maternally derived substances in a gestational age-dependent manner [17]. In addition, the biodistribution of cytokines and chemokines within gestational tissues is modulated by the actions of placental ABC transporters and, as such, may be involved in the pathogenesis of PTB.

Importantly, cultured human primary villous trophoblast cells exposed to LPS and to the viral double-stranded RNA (dsRNA) analogue polyinosinic:polycytidylic acid (polyI:C) exhibit markers of insulin resistance and increased amino acid uptake [18]. Treatment of trophoblast cells with cytokines, such as interleukin (IL)-1β and IL-6, elicited similar responses [19,20] indicating that infection may alter placental nutrient uptake and foetal transfer. However, the effect of infection on placental lipid uptake and foetal transfer is less well understood.

In the present study, we determined whether a bacterial infection at different stages of gestation modulates the levels of selected ABC transporters in the placenta, which in turn might alter foetal exposure to potentially harmful substrates. Furthermore, we investigated the maternal plasma and placental levels of lipid fractions and the mRNA expression of key placental lipid transporters in dams exposed to a sublethal LPS dose to elucidate the possible effects of LPS on altering lipid homeostasis related or unrelated to placental ABC transporter-mediated lipid exchange. We hypothesized that LPS exposure modifies the expression of key placental ABC and lipid transporters, as well as maternal and placental lipid homeostasis in a gestational age-dependent manner.

## Methods

### Animal experiments and study design

This study was approved by the Animal Care Committee of the Health Sciences Center, Federal University of Rio de Janeiro (CEUA-190/13) and registered within the Brazilian National Council for Animal Experimentation Control. The study complied with the “Principles of Laboratory Animal Care” generated by the National Society for Medical Research and the U.S. National Academy of Sciences Guide for the Care and Use of Laboratory Animals.

Virgin female and male C57BL/6 mice (8-10 weeks of age) were housed in a temperature-controlled room (23 °C) on a 12/12 h light/dark cycle, with free access to fresh food and water. Female mice in the oestrous phase (identified by vaginal cytology) were time-mated with C57BL/6 males and assigned (gestational day (GD) 0.5) to different groups. LPS (Sigma, Escherichia coli 055: B5; 150 μg/kg; intraperitoneal injection, i.p., in a single dose) or vehicle (i.p. injection of a single dose) was administered to mice in mid- (GD14.5/15.5) or late- (GD17.5/18.5) pregnancy for 24 or 4 h, respectively. Animals were euthanized at GD15.5 or GD18.5 at the end of the 4 or 24 h treatments with a single dose of LPS/vehicle, and maternal blood, foetuses and placentae were collected. Supplementary Figure 1 summarizes the design of the study.

Foetal and placental tissues were weighed, and three placentae per litter were selected for further study. The selection was based on placentae with weights closest to the mean, an approach that we and others have used previously [5,21–24]. The LPS dose (150 μg/kg) was selected because it has previously been shown to cause an acute maternal inflammatory response with less than 50% foetal death in mid-pregnancy at 4 h after treatment [25].

### qPCR

Total placental RNA was extracted using TRIzol reagent according to the manufacturer’s instructions (Life Technologies, CA, USA). The total RNA concentration was assessed using a nanophotometer (Implen, Munchen, Germany), and samples with RNA purity (260/280 absorbance) ratios ranging between 1.8 and 2.0 and with proven RNA integrity (confirmed through gel electrophoresis) were included in the study. Total RNA (1 μg) was reverse transcribed into cDNAs using the High Capacity cDNA Reverse Transcription Kit (Applied Biosystems, São Paulo, Brazil) according to the manufacturer’s instructions.

The mRNA levels of selected ABC transporters, lipid metabolism-related genes, proinflammatory cytokines and chemokines (Supplementary Table 1) were evaluated using qPCR according to the manufacturer’s recommendations (EVAGREEN; Solis Byodine, EUA) and using the Master Cycler Realplex system (Eppendorf, Germany) with the following cycling conditions: combined initial denaturation steps at 50°C (2 min) and 95°C (10 min), followed by 40 cycles of denaturation at 95°C (15 s), annealing at 60°C (30 s) and extension at 72°C (45 s). Relative gene expression was quantified using the 2-ΔΔCq method [26]. Assays with 95-105% efficiency were considered acceptable.

Gene expression was normalized to the geometric mean of selected reference genes in each experimental group, which exhibited stable expression levels following LPS challenge (Supplementary Table 1). The geometric mean expression of B2m and β-actin genes was used to normalize mRNA expression at GD15.5, whereas the geometric mean expression of Gapdh and Ywhaz reference genes was used to normalize mRNA expression at GD18.5. Intron-spanning primers, reverse transcriptase-negative samples and a melting curve analysis obtained for each qPCR were used to exclude DNA contamination.

### Histological and immunohistochemical staining

Placental discs were fixed with 4% buffered paraformaldehyde (PFA), dehydrated with increasing concentrations of ethanol, diaphanized in xylene, embedded in paraffin and sectioned (5 μm) using a Rotatory Microtome CUT 5062 (Slee Medical GmbH, Germany) for periodic acid-Schiff (PAS) staining and immunohistochemistry as previously described [5].

Briefly, PAS staining was performed by oxidizing placental sections with 0.5% periodic acid (Sigma-Aldrich, Missouri, USA) for 15 min. Sections were washed with distilled water and incubated (10 min at room temperature) with Schiff’s reagent (Merck, Germany), followed by haematoxylin (Proquímios, Rio de Janeiro, Brazil) staining. The junctional zone (Jz) interface between the maternal and foetal placental cellular components was visually identified, and the area of each region of interest, the Jz and the labyrinth zone (Lz), was measured using ImageJ software (National Institutes of Health, Maryland, USA).

Immunohistochemical staining was performed by incubating sections with Tris-EDTA buffer (pH 9.0) for 15 min, followed by immersion in sodium citrate buffer (pH 6.0) for 8 min (in a microwave). Sections were immersed in a bovine serum albumin (BSA3% - in PBS) solution to block nonspecific antibody binding sites and then incubated with primary antibodies against P-glycoprotein (P-gp-1:500; Santa Cruz Biotechnology, Texas, USA), breast cancer resistance protein (Bcrp-1:100; Merck Millipore, Massachusetts, USA) or Abcg1 (1:200; Abcam Plc, UK) overnight at 4 °C. BSA (3% - in PBS) solution was incubated with negative control sections instead of primary antibodies. Sections were then washed with PBS (3 × 5 min) and incubated with a biotin-conjugated secondary antibody (SPD-060 - Spring Bioscience, California, USA) for 1 h followed by an incubation for 1 h with streptavidin (SPD-060 - Spring Bioscience, California, USA). The reaction was halted with 3,3-diaminobenzidine (DAB) (SPD-060 - Spring Bioscience, California, USA) followed by hematoxylin (Proquímios, Brazil) staining.

Digital images of histological staining were acquired using a high-resolution Olympus DP72 camera (Olympus Corporation, Japan) attached to the Olympus BX53 microscope (Olympus Corporation, Japan). P-gp, Bcrp and Abcg1 staining were quantified using Image-Pro Plus 5.0 software (Media Cybernetics, Maryland, USA), where the percentage of stained tissue area was calculated and negative spaces were excluded. In each experimental group, 30 digital images per placenta (15 digital images for each labyrinthine and spongiotrophoblast area) were evaluated [5].

### Lipid analysis

Lipid extraction and analysis were performed using the method described by Bligh and Dyer with some modifications [27]. Maternal plasma (30 μl) and placentae (10 mg) were separately mixed with a solution containing chloroform (1 mL), methanol (2 mL) and water (0.8 mL) with intermittent shaking. After 2 h, the solution was centrifuged (1500 g, 20 min, 4 °C; Sorvall RC-5b; Sorvall Centrifuge, Newtown, CT, USA), the supernatant was collected, and a water-chloroform solution (1:1 v/v) was added. The mixture was shaken and centrifuged (1500 g, 20 min, 4°C), and the organic phase was removed and dried under nitrogen gas.

The lipid classes were separated by one-dimensional thin layer chromatography (TLC) for neutral lipids using a solution containing hexane, diethyl ether and acetic acid (60:40:1 v/v). Plates were immersed in a solution composed of 3% CuSO4 and 8% H3PO4 (v/v; 10 s) and then heated (110°C, 10 min) [28]. TLC plates were analyzed by densitometry using ImageMaster Total Lab 4.1 software (Total Lab Ltd, Newcastle, UK). Standards for each lipid species were used to identify different lipid classes (Sigma-Aldrich, Sao Paulo, Brazil). Their IDs and catalogue numbers are as follows: triacylglycerol, 1,3-dipalmitoyl-2-oleoylglycerol, #D2157; cholesteryl ester, cholesteryl oleate, #C9253; free fatty acids, palmitic acid, #76119; cholesterol, #C8667; and monoacylglycerol, #M2140.

A TLC plate with a standard curve for each of the lipid species was developed to quantify the lipid classes. Densitometric units from the standard curve were compared with densitometric units from the samples (which were already normalized to the unit of tissue or amount of plasma). TLC plates were developed in the mobile phase for 1 h. Supplementary figures 2 and 3 show the TLC plates with plasma and placenta samples from different groups.

### Measurement of plasma cytokine and chemokine levels

Maternal plasma was collected by cardiac puncture, transferred into heparinized tubes on ice, centrifuged (1,077 g, 15 min) and frozen (−80 °C). The levels of interleukin (Il)-1β, Il- 6, monocyte chemoattractant protein-1 (Mcp-1/Ccl2) and chemokine (C-X-C motif) ligand 1 (Cxcl1) were assessed using the MILLIPLEX-MAP kit Cytokine/Chemokine Magnetic Bead Panel (Merck Millipore, USA) according to the manufacturer’s recommendations. Fluorescence intensity was detected using a Luminex 200™ system (Merck Millipore, Massachusetts, USA).

### Statistical Analysis

Normality tests were applied followed by Student’s t-test or the nonparametric Mann-Whitney test to compare two variables. Pregnancy parameters were evaluated using the mean value of placentae/foetuses in each litter and not individuals [22]. For qPCR, PAS/immunostaining, lipid fraction analysis and cytokine/chemokine measurements, three placentae with the closest weight to the mean placental weight within each litter were selected. Thus, “n” represents the number of litters [5,21–23,25]. Values for all data are presented as the means ± SEM. GraphPad Prism 6 software (GraphPad Software, Inc., San Diego, CA, USA) was used to conduct statistical analyses, and differences were considered significant when P<0.05.

## Results

### Acute sublethal effects of LPS on pregnancy outcome

The sublethal LPS treatment elicited different pregnancy outcomes that varied according to the time of exposure and gestational age. At GD15.5, LPS at 4 h induced the death of 26% of foetuses, whereas at 24 h, an 84% foetal death rate was observed. Conversely, the same LPS dosage induced only 1 and 2% foetal death rates at GD18.5 after 4 h and 24 h of LPS exposure, respectively. However, a sublethal LPS treatment for 24 h in late pregnancy (GD18.5) induced a 64% increase in early labour compared to controls, i.e., fourteen of twenty-two dams exhibited signs of labour within 24 h (table 1) and, therefore, were not included in the study. Importantly, the average gestation length for C57BL/6 mice is 19.25 days (range GD18-22) [5,29], whereas PTB in C57BL/6 mice may occur prior to GD18 [5]. Since we were unable to determine the precise birth time within the 24 h limit of LPS treatment (i.e., GD17.5 or GD18.5) in our cohort, we opted to designate this mode of labour as early rather than preterm as a precaution. Of importance, the high percentage of foetal death at GD15.5 and the induction of early labour at GD18.5 prevented us from conducting further placental analysis in the 24 h groups. Thus, all following analyses were performed in tissues from foetuses that did not show signs of death and from dams that did not undergo early labour.

**Table 1.** Pregnancy outcomes following sublethal LPS challenge in mid and late pregnancy.

| LPS Injection Day | Group s | Exposure (h) | N (Dams) | Early Labor % | Foetal Death% | Foetal Weight (mg) | Placental Weight (mg) | F:P Ratio |
| --- | --- | --- | --- | --- | --- | --- | --- | --- |
| **E15.5** | Contro<br>l | 4 | 8 | 0 | 10 (8/81) | 410±30 | 95±2 | 4.3±0.35 |
|  | LPS |  | 8 | 0 | 36 (19/53) | 400±30 | 90±3 | 4.3±0.12 |
| **E14.5** | Contro<br>l | 24 | 6 | 0 | 3 (1/30) | ----- | ----- | ----- |
|  | LPS |  | 4 | 0 | 87 (20/23) | ----- | ----- | ----- |
| **E18.5** | Contro<br>l | 4 | 9 | 0 | 0 (0/35) | 1149±50 | 80±0.9 | 14.16±0.6 |
|  | LPS |  | 10 | 0 | 1 (1/68) | 1145±40 | 90*±2.0 | 12.80±0.64 |
| **E17.5** | Contro<br>l | 24 | 12 | 0 | 2.5 (2/80) | 1130±50 | 82±0.9 | 13.39±0.54 |
|  | LPS |  | 22 | 64 (14/22) | 2.0 (1/51) | 950*±60 | 92*±3 | 10.72**±0.72 |

Fetal death %, fetal weight, placental and fetal:placental weight (F:P) ratio values are extracted from pregnancies that did not undergo early labor, or from fetuses that did not show signs of demise. T-student test. \* $P < 0.05$ ; \*\* $P < 0.01$ .

### Maternal and placental pro-inflammatory/morphological response to acute sublethal LPS exposure

Maternal plasma levels of cytokines/chemokines associated with the pathogenesis of PTB, including Il-1β, Il-6, chemokine (C-X-C motif) ligand 1 (Cxcl1) and monocyte chemoattractant protein-1 (Mcp-1/Ccl2) [30,31] were assessed in dams following sublethal LPS exposure (4 h) at GD15.5 and GD18.5. At GD15.5, maternal plasma levels of Il-1β (P<0.01), Il- 6 (P<0.01), Cxcl1 (P<0.0001) and Ccl2 (P<0.01) were increased compared to the control group (Figure 1A–D). At GD18.5, we observed increased maternal plasma Il-6 (P<0.001), Cxcl1 (P<0.01) and Ccl2 levels (P<0.001) (Figure 1F–H), whereas Il-1β levels remained unchanged (Figure 1E).

**Figure 1.**
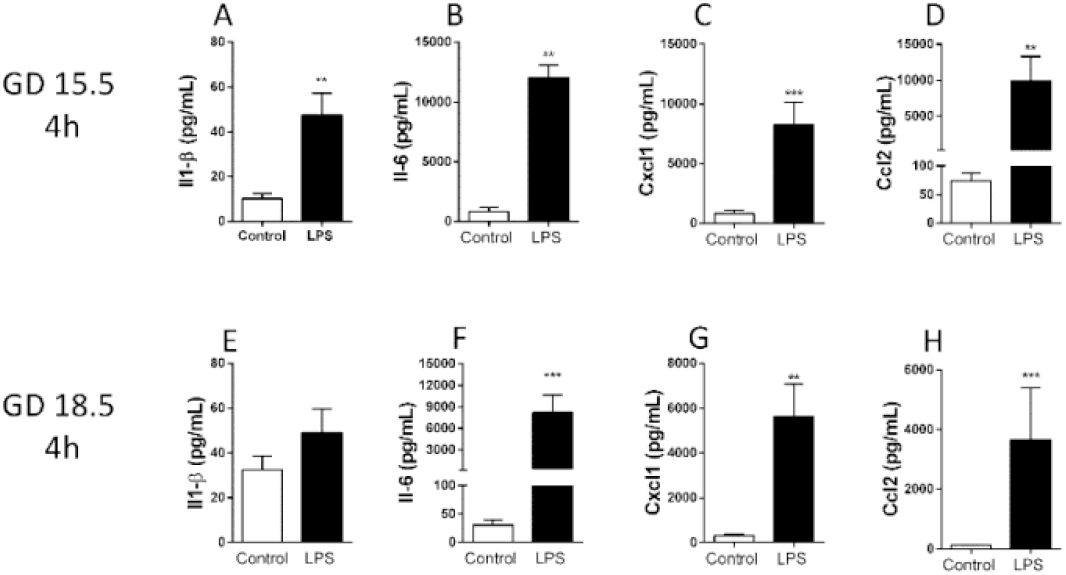
LPS challenge (4 h) elicited an acute maternal inflammatory response at GD 15.5 and 18.5. Maternal plasma levels of Il-1β (A and E), Il-6 (B and F), Cxcl1 (C and G) and Ccl2 (D and H) in dams at GD15.5 (control group, n=8, LPS group, n=8) and GD18.5 (control group, n=9, LPS group, n=10), respectively. Statistical analysis: unpaired Student’s t- test (GD15.5 and 18.5: Il1-β and Ccl2) or Mann-Whitney nonparametric test (GD15.5: Il-6 and Cxcl1; GD 18.5: Il-6). ** P<0.01, *** P<0.001.

In the placenta, the expression of the Il-6 and Cxcl1 (P<0.05) mRNA was significantly increased following LPS challenge at GD 15.5 and GD 18.5 (Figure 2). In contrast, Ccl2 mRNA expression was unchanged at GD15.5, but increased at GD18.5 (P<0.0001- Figure 2B).

**Figure 2.**
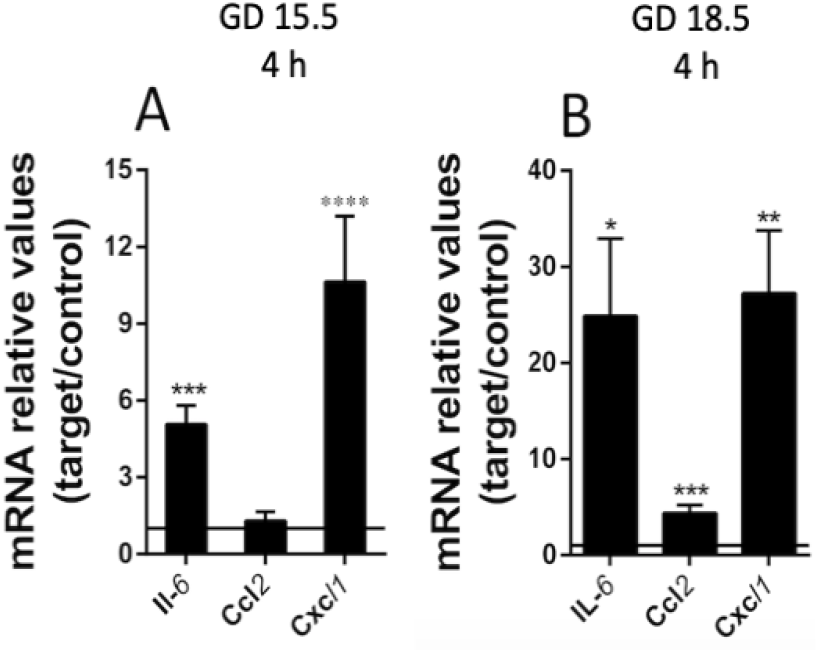
LPS insult (4 h) elicited an increase in selected placental cytokine/chemokine levels at GD 15.5 and 18.5. mRNA levels of placental Il6, Ccl2 and Cxcl1 at GD 15.5 (A - control group, n=8, LPS group, n=8) and GD 18.5 (B - control group, n=9, LPS group, n=10), 4 h after LPS exposure. Gene expression was normalized to the levels of the reference genes B2m and βactin (A), or Gapdh and Ywhaz (B). Statistical analysis: GD 15.5: Student’s t-test and GD 18.5: Student’s t -test (Lpl, Fabppm, Fatp1, Pparg) and Mann-Whitney nonparametric test (Cd36). * P< 0,05; ** P<0.01, *** P<0.001 **** P< 0.0001.

Placental weight was decreased 4 h after LPS exposure at GD18.5, compared to controls (Table 1). This led us to investigate whether LPS would elicit changes in the placental proportions of Lz and Jz. No differences in gross placental morphology, including changes in Lz and Jz areas, were identified in any of the groups investigated (Figure 3).

**Figure 3.**
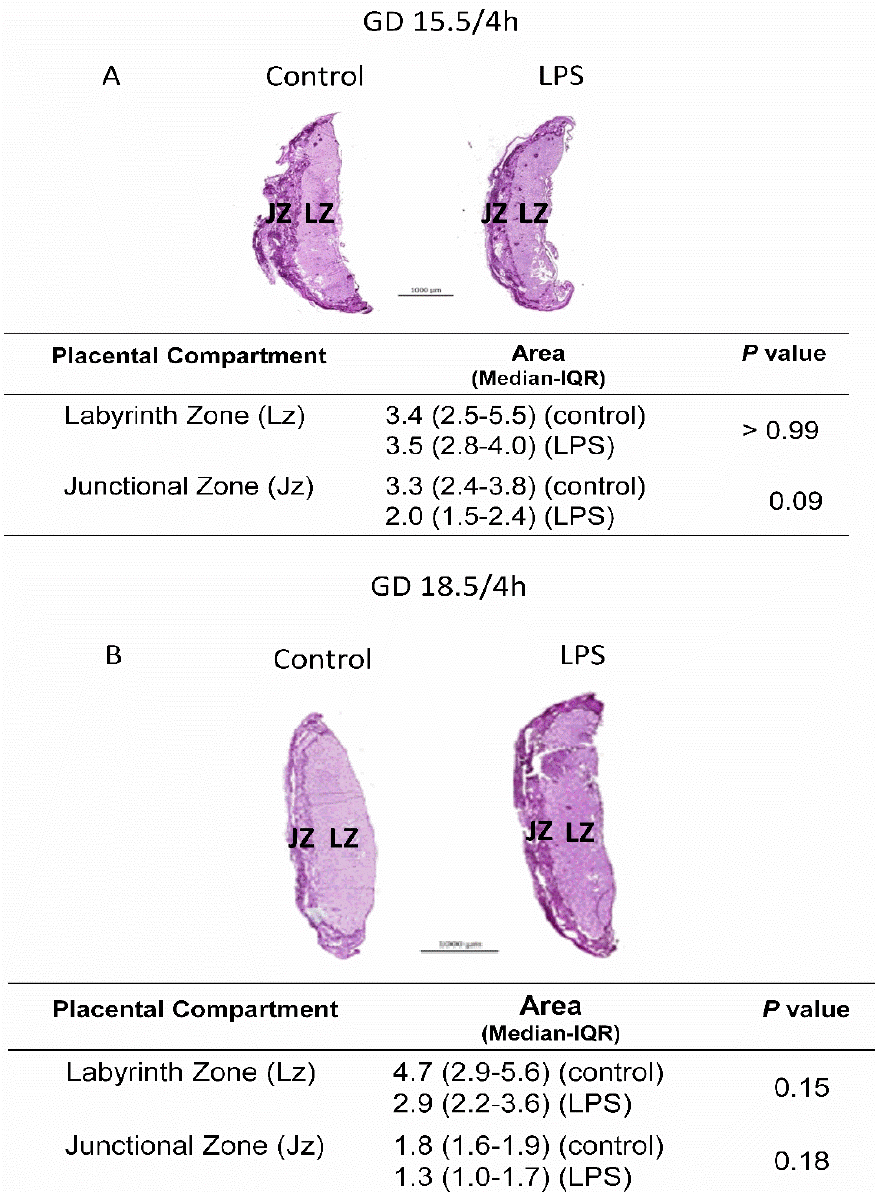
LPS (4 h) does not alter placental gross morphology. Analysis in the placental proportions of labyrinth (Lz) and junctional zone (Jz) of dams at GD15.5 (A) and 18.5 (B), 4 h after LPS exposure. Periodic acid-Schiff staining of con-trol and LPS-treated mice evaluating Lz and Jz areas. n=5/group. Statistical analysis: Mann-Whitney nonparametric test.

### Gestational age-dependent sublethal LPS effects on placental ABC transporters

To investigate how bacterial infection impacts placental efflux transport potential, we investigated the expression of the of Abca1, Abcb1a, Abcb1b, Abcb4, Abcc2, Abcc5, Abcf2, Abcg1 and Abcg2 mRNAs in the mouse placenta at GD15.5 or GD18.5 following LPS challenge (4 h). These specific ABC transporter genes were selected based on evidence showing their sensitivity to infection in other models and/or based on their importance to placental and yolk sac barrier function [5,14,15,32,33]. We did not observe significant differences after LPS treatments (4 h) at GD15.5 and GD18.5 (Figure 4).

**Figure 4.**
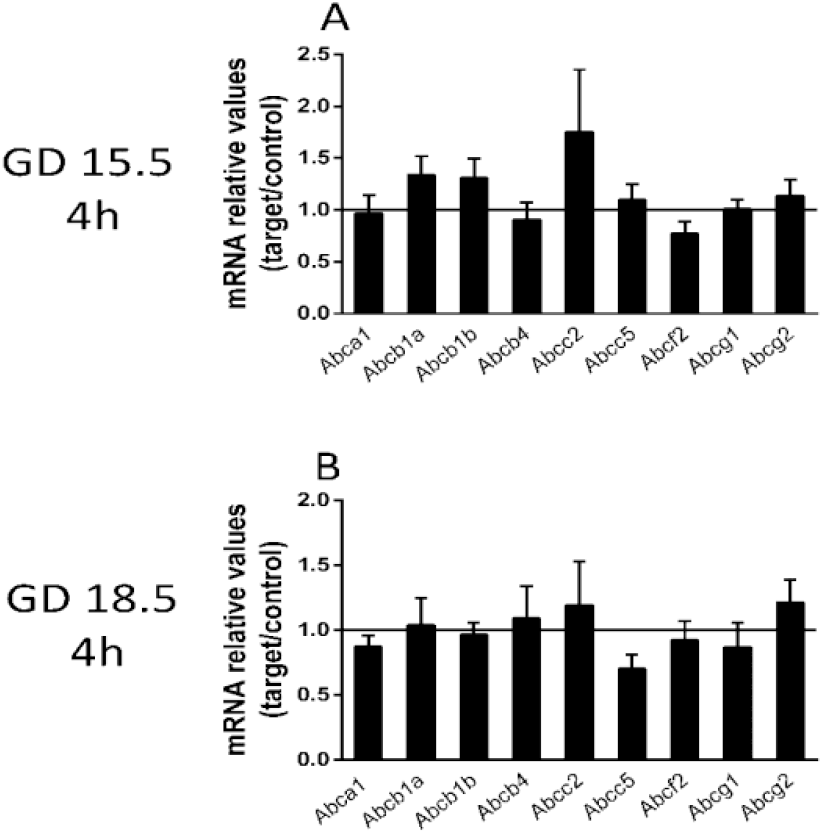
Sublethal LPS challenge (4 h) does not alter the expression of key ABC transporter genes at GDs 15.5 and 18.5. (A) Placental levels of the Abca1, Abcb1a, Abcb1b, Abcb4, Abcc2, Abcc5, Abcf2, and Abcg2 mRNA at GD15.5 (A) and at GD18.5 (B), 4 h after sublethal LPS administration. 15.5/4 h: n=8 (control group); n=8 (LPS group). 18.5/4 h: n=9 (control group)/n=10 (LPS group). Gene expression was normalized to the levels of the reference genes (A) B2m and βactin or (B) Gapdh and Ywhaz. Statistical analysis: GD 15.5: Student’s t test and GD 18.5: Student’s t-test (Abcb1b, Abcb4, Abcc5, Abcf2, Abcg1) and Mann-Whitney nonparametric test (Abca1, Abcb1a, Abcc2, Abcg2).

P-gp and Bcrp are key multidrug resistance transporters that have been shown to play roles in foetal protection, whereas the lipid transporter Abcg1 is important for foetal lipid transfer. Therefore, these three ABC transporters were specifically chosen for further examination. P-gp immunostaining was detected in the cellular membrane of labyrinthine cells, with variable staining in the cellular membrane and cytoplasm of spongiotrophoblast cells (Figure 5). Bcrp exhibited a similar placental distribution pattern, but with faint and variable nuclear staining in labyrinthine cells. Generally, greater Bcrp cytoplasmic labelling was observed in labyrinthine and spongiotrophoblast cells compared to P-gp-labelled placental cells (Figure 6). The lipid transporter Abcg1 was predominantly localized to cellular membranes in the Lz and Jz, with some variable cytoplasmic staining throughout these layers (Figure 7). Supplementary figures 4, 5 and 6 depict higher magnification photomicrographies of placental P-gp, Bcrp and Abcg1 staining, respectively.

**Figure 5.**
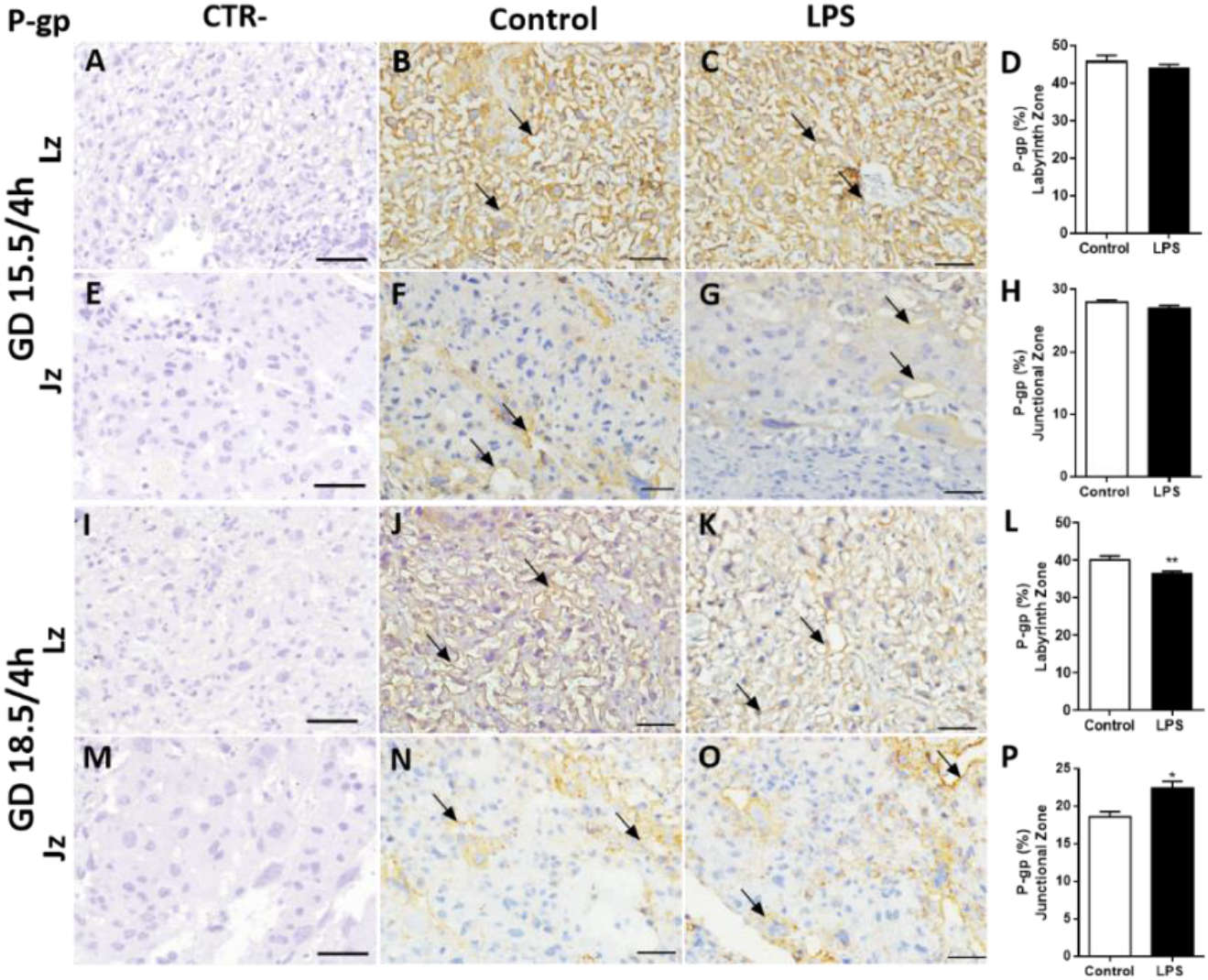
LPS challenge modulates P-gp protein in the mouse placenta. Representative photomicrographies of im-munohistochemistry staining (arrows) and semiquantitative analysis of P-gp in the labyrinth zone (Lz) and junctional zone (Jz) at GD 15.5/4h (A-H) and GD 18.5/4h (I-P) after LPS administration (4 h). Graphs represent the % of positively stained cells. n=5/group. Scale bar = 50 μm). Statistical analysis: Student’s t-test. * P< 0,05; ** P<0.01 * P< 0,05; ** P<0.01.

**Figure 6.**
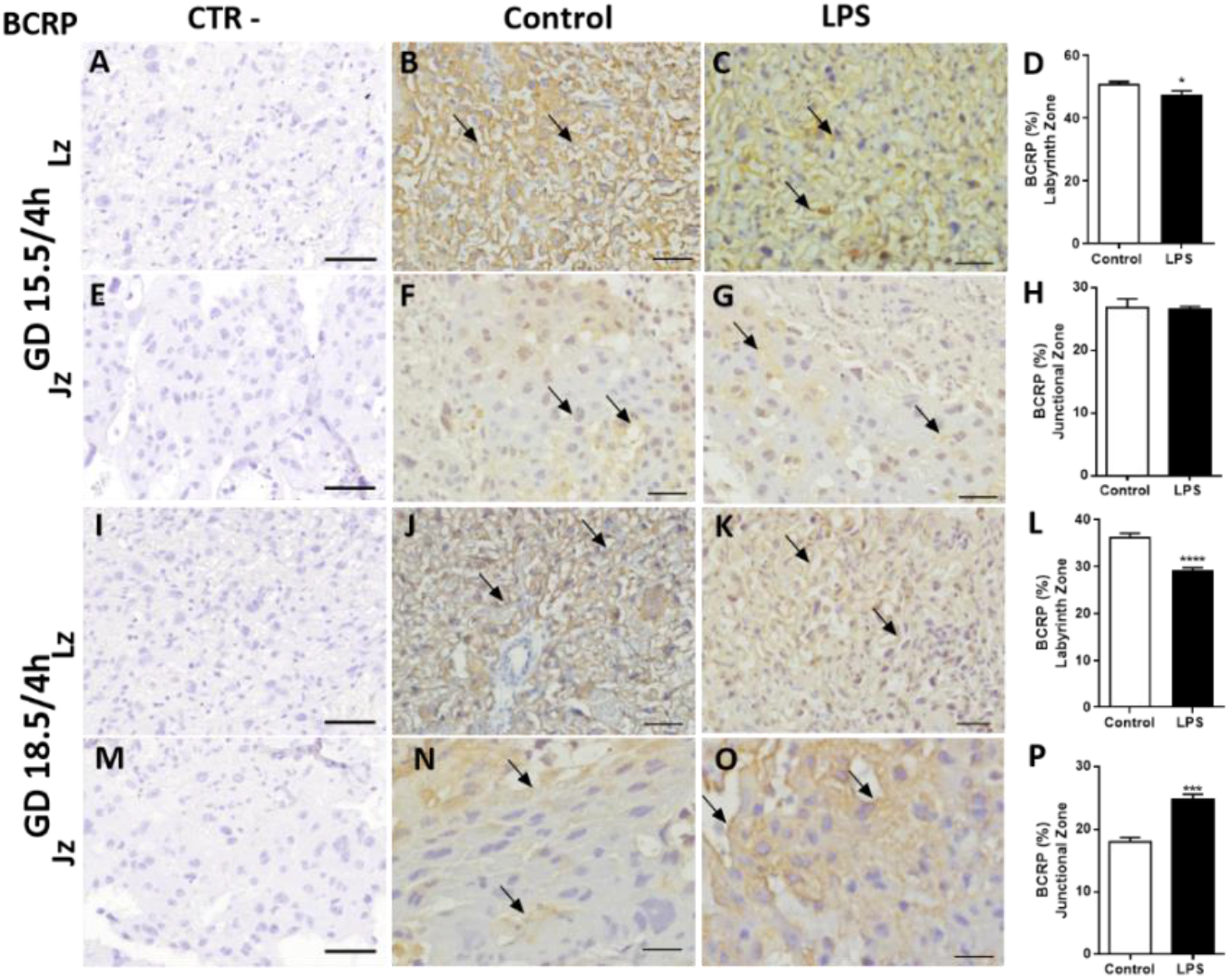
LPS challenge modulates BCRP protein in the mouse placenta. Representative photomicrographies of immunohistochemistry staining (arrows) and semiquantitative analysis of BCRP in the labyrinth zone (Lz) and junctional zone (Jz) at GD 15.5/4h (A-H) and GD 18.5/4h (I-P) after LPS administration (4 h). Graphs represent the % of positively stained cells. n=5/group. Scale bar = 50 μm). Statistical analysis: Student’s t-test. * P< 0,05; *** P<0.001 **** P< 0.0001.

**Figure 7.**
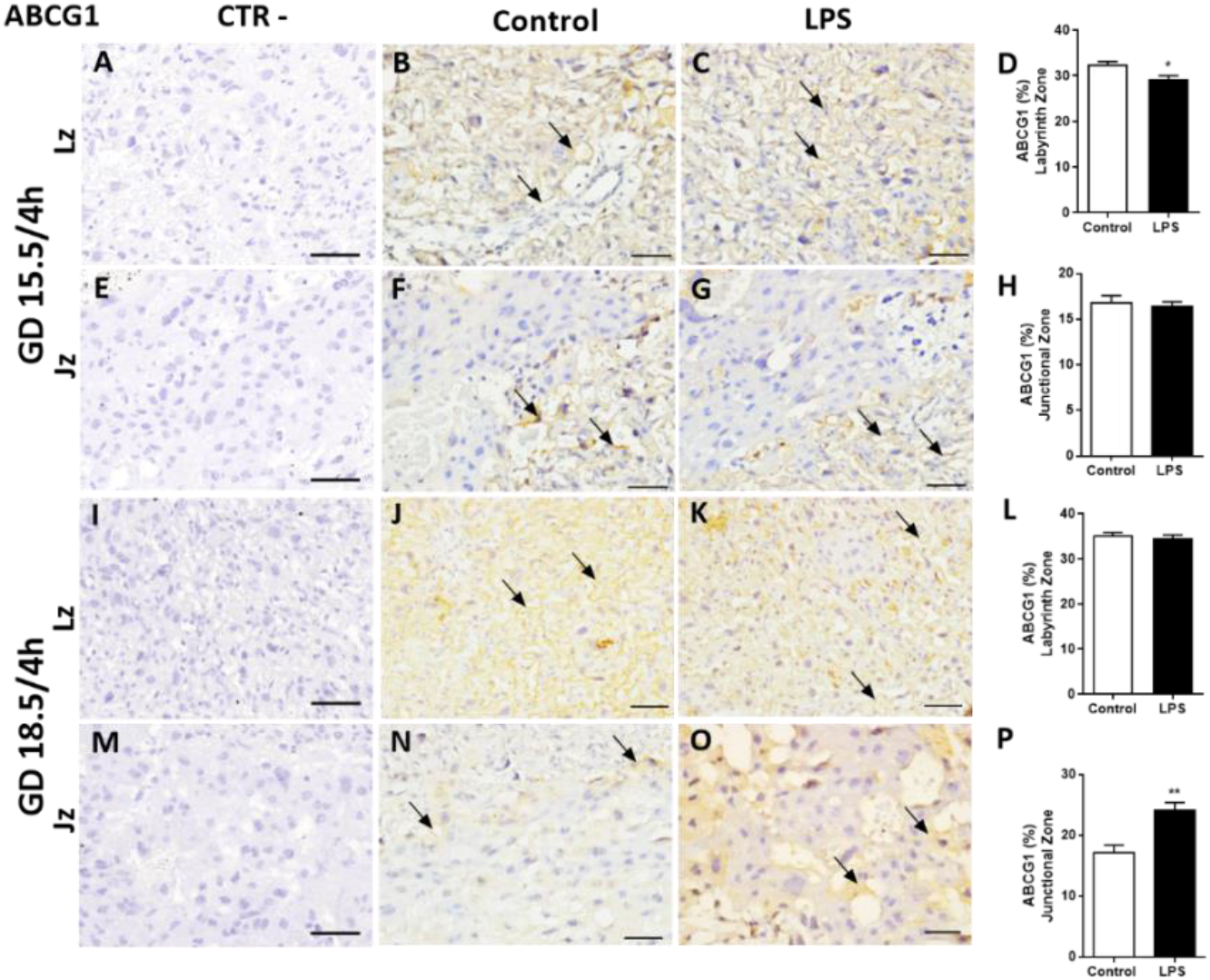
LPS challenge modulates Abcg1 protein in the mouse placenta. Representative photomicrographies of immunohistochemistry staining and semiquantitative analysis of Abcg1 in the labyrinth zone (Lz) and junctional zone (Jz) at GD 15.5/4h (A-H), GD 18.5/4h (I-P) after LPS administration (4 h). Graphs represent the % of positively stained cells. n=5/group. Scale bar = 50 μm). Statistical analysis: Student’s t-test. * P< 0,05; ** P<0.01.

Using a semiquantitative analysis, we identified reduced P-gp staining intensity in the Lz at GD18.5 (P<0.01) after LPS administration (4 h), compared to controls (Figure 5L). No differences in Lz-P-gp staining were detected after LPS treatment at GD15.5 (Figure 5D). The Bcrp staining intensity in the Lz was reduced at GD15.5 (P<0.05) and at GD 18.5 (P<0.0001) (Figure 6D and L). A lower Abcg1 staining intensity was observed in labyrinth cells at GD 15.5 (P<0.05), whereas at GD18.5, it remained unchanged (Figure 7D and L). In contrast, P-gp, Bcrp and Abcg1 intensity in spongiotrophoblast cells were higher at GD18.5 (P<0.05) 4h after LPS exposure (Figures 5P, 6P and 7P respectively). No differences in the levels of P-gp, Bcrp and Abcg1 in spongiotrophoblast cells were identified at GD15.5 after LPS treatment (Figure 5,6 and 7H, respectively).

### LPS alters maternal and placental lipid homeostasis throughout pregnancy

Since we observed an inhibitory effect of LPS on Abcg1 expression (a lipid transporter) in the Lz, we investigated the impact of sublethal LPS exposure (4 h) on the maternal plasma and placental levels of various lipid classes (triacylglycerol, free fatty acids, cholesterol ester, cholesterol, monoacylglycerol, and phospholipids) at GDs 15.5 and 18.5.

We observed significant alterations in placental lipid levels following sublethal LPS challenge. Triacylglycerol, free fatty acid, cholesterol ester and free cholesterol levels were decreased at GD15.5 compared to the control groups (P<0.01, Table 2), whereas triacylglycerol, free fatty acid, free cholesterol, monoacylglycerol and phospholipid levels were increased at GD18.5 compared to the control groups (P<0.05, Table 3). The levels of the lipid classes monoacylglycerol and phospholipid remained unchanged at GD15.5 (Table 2), whereas cholesterol ester levels did not exhibit alterations at GD18.5 (Table 3).

**Table 2.** LPS challenge modulates placental lipid fractions at GD15.5 after LPS administration.

| Placental lipid fractions | Groups | mean ± SEM<br>($\mu\text{g}/\text{mg}$) | P value |
| --- | --- | --- | --- |
| Triacylglycerol | Control<br>LPS | 0.05±0.002<br>0.03±0.002 | 0.0001*** |
| Free Fatty Acids | Control<br>LPS | 0.014±0.001<br>0.008±0.001 | 0.0043** |
| Cholesterol Ester | Control<br>LPS | 0.18±0.02<br>0.10±0.01 | 0.005** |
| Cholesterol | Control<br>LPS | 0.05±0.003<br>0.03±0.004 | 0.0043** |
| Monoacylglycerol | Control<br>LPS | 0.02±0.0002<br>0.02±0.002 | 0.66 |
| Phospholipids | Control<br>LPS | 0.073±0.003<br>0.077±0.005 | 0.51 |

15.5/4h: n=6 (control group); n=5 (LPS group). Statistical analysis: Student's t-test. \*\* P<0.01, \*\*\* P<0.001.

**Table 3:** LPS challenge modulates placental lipid fractions at GD18.5 after LPS administration.

| Placental lipid fractions | Groups | mean ± SEM<br>($\mu\text{g}/\text{mg}$) | P value |
| --- | --- | --- | --- |
| Triacylglycerol | Control<br>LPS | 0.04±0.009<br>0.08±0.008 | 0.03* |
| Free Fatty Acids | Control<br>LPS | 0.004±0.0005<br>0.03±0.001 | <0.0001**** |
| Cholesterol Ester | Control<br>LPS | 0.23±0.02<br>0.19±0.01 | 0.09 |
| Cholesterol | Control<br>LPS | 0.026±0.001<br>0.04±0.0008 | <0.0001**** |
| Monoacylglycerol | Control<br>LPS | 0.03±0.002<br>0.04±0.003 | 0.0002*** |
| Phospholipids | Control<br>LPS | 0.065±0.002<br>0.071±0.002 | 0.04* |

18.5/4 h: n=9 (control group); n=7 (LPS group). Statistical analysis: Student's t-test. \* P<0,05, \*\*\* P<0.001, \*\*\*\* P<0.0001.

We detected increased triacylglycerol levels and decreased free fatty acids and cholesterol ester levels in the maternal plasma from the LPS-exposed group at GD15.5 compared to the control group (P<0.001, table 4). At GD18.5, the triacylglycerol levels were decreased in the LPS group compared to the control group (P<0.05, table 5). No differences were observed in the levels of free cholesterol, monoacylglycerol and phospholipid classes at either gestational age, whereas the cholesterol ester contents remained unchanged at GD18.5 (tables 4 and 5).

**Table 4.** LPS challenge modulates plasma lipid fractions at GD15.5 after LPS administration.

| Plasma lipid fractions | Groups | mean ± SEM<br>($\mu\text{g}/\mu\text{l}$) | P value |
| --- | --- | --- | --- |
| Triacylglycerol | Control<br>LPS | 0.31±0.02<br>0.56±0.02 | <0.0001**** |
| Free Fatty Acids | Control<br>LPS | 0.14±0.01<br>0.06±0.003 | <0.0001**** |
| Cholesterol Ester | Control<br>LPS | 0.31±0.02<br>0.19±0.01 | 0.0003*** |
| Cholesterol | Control<br>LPS | 0.11±0.02<br>0.13±0.02 | 0.59 |
| Monoacylglycerol | Control<br>LPS | 0.04±0.003<br>0.06±0.006 | 0.095 |
| Phospholipids | Control<br>LPS | 0.30±0.07<br>0.38±0.03 | 0.26 |

15.5/4 h: n=7 (control group); n=8 (LPS group). Statistical analysis: Student's t-test. \*\*\* P<0.001 \*\*\*\* P<0.0001.

**Table 5:** LPS challenge modulates plasma lipid fractions at GD15.5 after LPS administration.

| Plasma lipid fractions | Groups | mean ± SEM<br>($\mu\text{g}/\mu\text{l}$) | P value |
| --- | --- | --- | --- |
| Triacylglycerol | Control<br>LPS | 0.76±0.07<br>0.57±0.03 | 0.04* |
| Free Fatty Acids | Control<br>LPS | 0.06±0.01<br>0.08±0.01 | 0.44 |
| Cholesterol Ester | Control<br>LPS | 0.16±0.03<br>0.22±0.07 | 0.4 |
| Cholesterol | Control<br>LPS | 0.12±0.02<br>0.15±0.02 | 0.23 |
| Monoacylglycerol | Control<br>LPS | 0.07±0.009<br>0.06±0.007 | 0.26 |
| Phospholipids | Control<br>LPS | 0.19±0.01<br>0.20±0.008 | 0.19 |

18.5/4h: n=9 (control group); n=7 (LPS group). Statistical analysis: Student's t-test. \* P<0,05.

Placental mRNA expression of key lipid transporters and lipid metabolism-related genes following sublethal LPS treatments was assessed. At GD15.5 (4 h), LPS exposure resulted in lower fatty acid binding protein associated with plasma membrane (Fabppm) mRNA levels (p<0.05), whereas at GD18.5 (4 h), increased placental Fat/Cd36 translocase levels (p<0.01) were detected (Figure 8A–B). The mRNA expression of other lipid transporters and metabolism-related genes, such as Pparg, fatty acid transporter family protein (Fatp1) and lipase lipoprotein (Lpl), remained unchanged throughout pregnancy.

**Figure 8.**
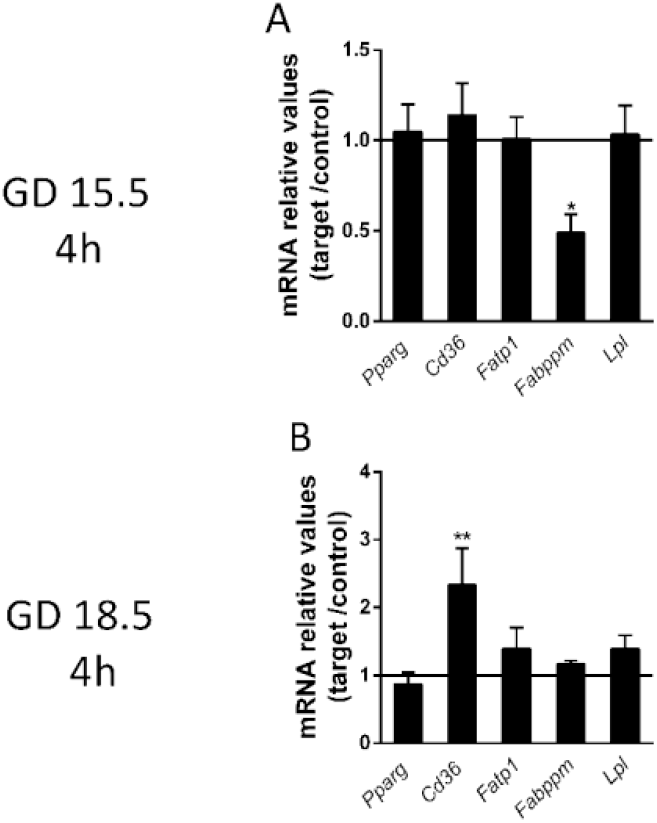
LPS challenge modulates the mRNA expression of lipid transporters and associated proteins at GDs 15.5 and 18.5, 4 h after the insult. (A-B) Quantification of the relative mRNA expression of selected lipid transporters (Pparg and Cd36) and associated proteins (Fatp1, Fabppm and Lpl) at GD 15.5 and 18.5 4 h after LPS challenge. 15.5/4 h n=8 (control group); n=8 (LPS group). 18.5/4 h: n=9 (control group)/n=10 (LPS group). Gene expression was normalized to the levels of the reference genes (A) B2m and actin, or (B) Gapdh and Ywhaz. Student’s t test. Statistical analysis: GD 15.5: Stu-dent’s t-test and GD 18.5: Student’s t-test (Lpl, Fabppm, Fatp1, Pparg) and Mann-Whitney nonparametric test (Cd36). * P< 0,05; ** P<0.01.

## Discussion

We have documented the effects of bacterial infection on placental ABC transporters and lipid homeostasis. LPS caused substantial foetal death at GD15.5 and triggered early labour in late pregnancy (GD18.5) without inducing foetal demise. These effects were likely mediated by distinct maternal and placental cytokine/chemokine responses to LPS throughout pregnancy and were associated with distinct expression profiles of placental efflux and lipid transporters, as well as changes in maternal and placental lipid levels.

Our present findings showing a greater susceptibility to LPS at GD15.5 are consistent with previous studies from our group [25]. However, here, we also show that sublethal LPS challenge in late pregnancy increased early labour, probably by intensifying the maternal and placental output of labour-inducing cytokines and chemokines. Our data are consistent with one study conducted in rats showing that LPS treatment (24 h) at GD18.5 induced early labour in 55% of pregnancies [34], a percentage similar to the value reported in our study (64%). Furthermore, LPS administration in late gestation decreased the foetal weight, stimulated placental growth and reduced the F:P weight ratio. These changes might indicate an impairment of placental nutrient transport efficiency that prevents foetuses from attaining optimal growth potential [30,31,34,35] and predisposing them to poor long-term health outcomes [35].

Acute sublethal LPS exposure during pregnancy elicited an intense maternal and placental proinflammatory response, which varied across pregnancy. We observed a marked increase in a number of key proinflammatory factors (IL-6 and IL-8 and CCL2) related to PTB. In contrast, plasma Il-1β levels and placental levels of the Ccl2 mRNA were not consistently affected by LPS, since they did exhibit a gestational age-dependent expression profile. IL-1β, IL-6 and IL-8 are commonly upregulated by infection and induce prostaglandin (PG)E2 and PGF2 production to stimulate myometrial contractility and PTB [31,36,37]. CCL2 is subsequently upregulated in the human myometrium during preterm and term labour and recruits infiltrating leukocytes into the myometrium to amplify local inflammation and trigger the onset of labour [38]. CCL2 also induces the expression of proinflammatory cytokines, prostaglandins and leukotrienes in the myometrium during late pregnancy [39,40]. Importantly, Tlr4 activation induces the myometrial production of Il-1β, Il-6 and Ccl2 via nuclear factor- κB (NF-κB) and p38 mitogen-activated protein kinase (MAPK) activation [41].

In the present study, LPS stimulated placental Ccl2 expression in a gestational time-dependent manner, suggesting that this cytokine may participate in the pathogenesis of infective chorioamnionitis and the induction of early labour. Furthermore, our data extracted from pregnancies that did not undergo early labour and at an earlier time point (4 h) indicate that a lack of Il- 1β upregulation in these pregnancies may have been important in preventing an earlier onset of labour. Future studies comparing the maternal and placental inflammatory responses of pregnancies that did and did not undergo LPS-induced early labour are required to test this hypothesis.

We evaluated the mRNA and protein expression/localization of selected ABC transporters involved in the biodisposition of clinically relevant substrates across pregnancy to understand how a sublethal bacterial infection alters placental efflux transport potential. As shown in our previous study, a sublethal LPS challenge (150 μg/kg) impairs placental P- gp activity at GD15.5 (4 h) in C57BL/6 mice, with no changes in placental Abcb1a and Abcb1b mRNA levels [25]. In the present study, no changes in P-gp-encoding genes (Abcb1a and Abcb1b) were observed at GD15.5 and GD18.5 (4 h), however, lower labyrinthine P-gp staining intensity at GD18.5 (4 h) was detected. Placental levels of P-gp and its encoding genes are developmentally regulated in both rodents and humans, i.e., expression decreases towards term [42,43], resulting in reduced foetal protection against P-gp substrates in late pregnancy. Based on our results, a late-term bacterial infection may further decrease this already limited P-gp-mediated barrier function. These changes will likely increase the levels of cytokines and chemokines within gestational tissues, causing foetal demise and morbidity and/or inducing preterm/early labour pathways.

The results from the present and previous studies suggest that bacterial infection has the potential to increase foetal accumulation of P-gp substrates (cytokines, chemokines, endogenous and synthetic glucocorticoids, antibiotics, antiretrovirals, etc.) [14] both in earlier stages of pregnancy due to impaired placental P-gp activity [25] and in later stages of pregnancy by decreasing labyrinthine-P-gp expression. These changes will likely increase the levels of cytokines and chemokines within gestational tissues, causing foetal demise and morbidity and/or inducing preterm/early labour pathways.

Consistent with our data, P-gp is downregulated in a) human first trimester placental explants exposed to LPS [16], b) human PTB placentae with chorioamnionitis [15] and c) placentae from pregnant malaria-infected C57BL/6 mice exhibiting high rates of PTB [5]. Furthermore, P-gp activity is impaired in the developing foetal blood brain barrier of pregnant C57BL/6 mouse exposed to PolyI:C, a TLR3 viral mimic [44]. Together, these studies suggest that lower placental Abcb1b and P-gp levels are associated with infection-driven PTB [5,15] and/or earlier labour and delivery. The effects of lower placental P-gp levels on foetal health and postnatal development require further investigation.

A lower level of Bcrp immunostaining was observed in the Lz at GDs 15.5 and 18.5, following LPS exposure. Placental Bcrp levels have been shown to be downregulated in human first trimester placental explants treated with LPS [16], in malaria-infected murine placentae [5] and in HTR8/SVneo (human extravillous trophoblast (EVT)-like) cells exposed to either LPS or to the TLR8 viral mimic single stranded RNA (ssRNA) [45]. Thus, different infective agents/challenges during pregnancy, including bacterial infection, have the potential to increase foetal accumulation of Bcrp substrates during pregnancy. However, BCRP was upregulated in human PTB placentae with chorioamnionitis [15] which is commonly induced by polymicrobial infection [7], and after treatment of trophoblastic cells with PGE2 [46], indicating that the nature of infective/inflammatory stimuli determines the trophoblastic-BCRP modulatory response.

Interestingly, the P-gp, Bcrp and Abcg1 staining intensity in the spongiotrophoblast was higher at GD18.5 following LPS (4 h) administration but not different at GD15.5. The role of ABC transporters in spongiotrophoblast cells is far less understood. Bcrp immunolocalization has been previously reported in spongiotrophoblasts, which remained unaltered in pregnancies in which the mother underwent nutritional manipulations [21]. Spongiotrophoblasts comprise the mouse placental Jz and provide structural support for the growth of the labyrinthine villi and limit foetal endothelium overgrowth [47]. However, very little is known about the possible functions of ABC transporters in the mouse Jz and how infection and inflammation impact Jz function.

Labyrinthine Abcg1 expression was also downregulated by LPS at GD15.5. Abcg1 is a cholesterol and phospholipid efflux transporter predominantly localized to the basolateral membrane of the syncytiotrophoblast and in the endothelium of the foetal capillaries of the human placenta, suggesting that it mediates lipid efflux from the maternal compartment to the foetal compartment [14]. However, the directionality of Abcg1-mediated lipid exchange in the mouse placenta has yet to be described, preventing us from postulating on the effects of sublethal LPS exposure on placental Abcg1-mediated biodisposition of cholesterol and other lipids.

Maternal plasma and placental lipid levels were subsequently investigated to better understand how a bacterial infection alters lipid homeostasis at the maternal-foetal interface. Maternal levels of triacylglycerol and free fatty acids were altered after LPS exposure at GD15.5. Similar results have been reported in hepatic tissue following LPS exposure [48], which occurs in a gestational time-dependent manner. In fact, relevant alterations in plasma lipoproteins occur during injury or infection [49] or in patients with sepsis who occasionally present with hypertriglyceridemia [49]. The higher plasma triacylglycerol levels may be directly related to the inflammatory status in pregnant mice and may be associated with the high rates of foetal death at GD15.5. Nevertheless, higher placental triacylglycerol and free fatty acids levels were detected at GD18.5. This later effect may be an attempt to circumvent the lower foetal weight and one possible mechanism responsible for the higher placental weight observed in this group.

Placental fatty acid transport is modulated by different transport systems, including fatty acid transporter family proteins (FATPs), fatty acid binding proteins (FABPs), fatty acid binding protein associated with plasma membrane (FABPpm), lipase lipoproteins (LPL) and Fat/cd36 translocase located on both apical and basolateral membranes of the trophoblast [50,51]. We observed decreased placental Fabppm at GD15.5 and increased Fat/cd36 mRNA expression at GD18.5 after LPS exposure (4 h). This finding may at least partially explain the different patterns of free fatty acid accumulation we observed in the present study. Foetuses (and intrauterine tissues) in pregnancies complicated by maternal bacterial infection may be exposed to higher levels of cytokines/chemokines, drugs and environmental toxins present in the maternal circulation. These changes may also be associated with suboptimal placental lipid storage. Combined, these changes may contribute to the inflammatory PTB pathways and lead to the onset of PTB or early labour.

In conclusion, this sublethal LPS model of bacterial infection during pregnancy may induce foetal death or early labour, depending on the gestational age. These outcomes are associated with specific maternal and placental inflammatory responses, altered expression of ABC and lipid transporters and altered maternal and placental lipid homeostasis.

## Acknowledgments

This study was supported by the Bill & Melinda Gates Foundation (MCTI/CNPq/MS/SCTIE/Decit/Bill and Melinda Gates 05/2013; OPP1107597), the Canadian Institutes for Health Research (SGM: Foundation-148368), Conselho Nacional de Desenvolvimento Científico e Tecnológico (CNPq; 304667/2016-1, 422441/2016-3, 303734/2012-4, 422410/2016-0); Coordenação de Aperfeiçoamento Pessoal de Nível Superior (CAPES, finance Code 001); Fundação de Amparo à Pesquisa do Estado do Rio de Janeiro (FAPERJ, CNE 2015/E26/203.190/2015); and PRPq-Universidade Federal de Minas Gerais (PRPq-UFMG, 26048).

## Competing Interests

The authors declare no competing interests.

## Notes

### Competing Interest Statement

The authors have declared no competing interest.

### Summary of Updates

A minor typo in table 2 was corrected.Figures 1 and 3 were corrected and 2 new supplementary figures were added.

## References

1. Harrison, M.S.; Goldenberg, R.L. Global burden of prematurity. Semin. Fetal Neonatal Med. 2016, 21, 74–79, doi:10.1016/j.siny.2015.12.007.

2. Goldenberg, R.L.; Culhane, J.F.; Iams, J.D.; Romero, R. Preterm Birth 1: Epidemiology and Causes of Preterm Birth. Obstet. Anesth. Dig. 2009, doi:10.1097/01.aoa.0000344666.82463.8d.

3. Bloise, E.; Torricelli, M.; Novembri, R.; Borges, L.E.; Carrarelli, P.; Reis, F.M.; Petraglia, F. Heat-killed lactobacillus rhamnosus GG modulates urocortin and cytokine release in primary trophoblast cells. Placenta 2010, 31, 867–872, doi:10.1016/j.placenta.2010.04.007.

4. Novembri, R.; Torricelli, M.; Bloise, E.; Conti, N.; Galeazzi, L.R.; Severi, F.M.; Petraglia, F. Effects of urocortin 2 and urocortin 3 on IL-10 and TNF-α; expression and secretion from human trophoblast explants. Placenta 2011, 32, 969–974, doi:10.1016/j.placenta.2011.09.013.

5. Fontes, K.N.; Reginatto, M.W.; Silva, N.L.; Andrade, C.B.V.; Bloise, F.F.; Monteiro, V.R.S.; Silva-Filho, J.L.; Imperio, G.E.; Pimentel-Coelho, P.M.; Pinheiro, A.A.S.; et al. Dysregulation of placental ABC transporters in a murine model of malaria-induced preterm labor. Sci. Rep. 2019, 9, 1–13, doi:10.1038/s41598-019-47865-3.

6. Challis, J.R.; Lockwood, C.J.; Myatt, L.; Norman, J.E.; Strauss, J.F.; Petraglia, F. Inflammation and pregnancy. Reprod. Sci. 2009, 16, 206–215, doi:10.1177/1933719108329095.

7. Conti, N.; Torricelli, M.; Voltolini, C.; Vannuccini, S.; Clifton, V.L.; Bloise, E.; Petraglia, F. Term histologic chorioamnionitis: A heterogeneous condition. Eur. J. Obstet. Gynecol. Reprod. Biol. 2015, 188, 34–38, doi:10.1016/j.ejogrb.2015.02.034.

8. Nadeau, H.C.G.; Subramaniam, A.; Andrews, W.W. Infection and preterm birth. Semin. Fetal Neonatal Med. 2016, 21, 100–105, doi:10.1016/j.siny.2015.12.008.

9. Beutler, B.; Hoebe, K.; Du, X.; Ulevitch, R.J. How we detect microbes and respond to them: the Toll-like receptors and their transducers. J. Leukoc. Biol. 2003, 74, 479–485, doi:10.1189/jlb.0203082.

10. Ogando, D.G.; Paz, D.; Cella, M.; Franchi, A.M. The fundamental role of increased production of nitric oxide in lipopolysaccharide-induced embryonic resorption in mice. Reproduction 2003, 125, 95–110, doi:10.1530/rep.0.1250095.

11. Leazer TM, Barbee B, Ebron-McCoy M, Henry-Sam GA, R.J. Role of the maternal acute phase response and tumor necrosis factor alpha in the developmental toxicity of lipopolysaccharide in the CD-1 mouse. Reprod Toxicol. 2002, 16, 173–9, doi:https://doi.org/10.1016/S0890-6238(02)00011-4.

12. Guo, Y.; Ma, Z.; Kou, H.; Sun, R.; Yang, H.; Smith, C.V.; Zheng, J.; Wang, H. Synergistic effects of pyrrolizidine alkaloids and lipopolysaccharide on preterm delivery and intrauterine fetal death in mice. Toxicol. Lett. 2013, 221, 212–218, doi:10.1016/j.toxlet.2013.06.238.

13. Elovitz, M.A.; Brown, A.G.; Breen, K.; Anton, L.; Maubert, M.; Burd, I. Intrauterine inflammation, insufficient to induce parturition, still evokes fetal and neonatal brain injury. Int. J. Dev. Neurosci. 2011, 29, 663–671, doi:10.1016/j.ijdevneu.2011.02.011.

14. Bloise, E.; Ortiga-Carvalho, T.M.; Reis, F.M.; Lye, S.J.; Gibb, W.; Matthews, S.G. ATP-binding cassette transporters in reproduction: A new frontier. Hum. Reprod. Update 2016, 22, 164–181, doi:10.1093/humupd/dmv049.

15. Do Imperio, G.E.; Bloise, E.; Javam, M.; Lye, P.; Constantinof, A.; Dunk, C.; Dos Reis, F.M.; Lye, S.J.; Gibb, W.; Ortiga-Carvalho, T.M.; et al. Chorioamnionitis induces a specific signature of placental ABC transporters associated with an increase of miR-331-5p in the human preterm placenta. Cell. Physiol. Biochem. 2018, 45, 591–604, doi:10.1159/000487100.

16. Lye, P.; Bloise, E.; Javam, M.; Gibb, W.; Lye, S.J.; Matthews, S.G. Impact of bacterial and viral challenge on multidrug resistance in first- and third-trimester human placenta. Am. J. Pathol. 2015, doi:10.1016/j.ajpath.2015.02.013.

17. Imperio, G.E.; Javam, M.; Lye, P.; Constantinof, A.; Dunk, C.E.; Reis, F.M.; Lye, S.J.; Gibb, W.; Matthews, S.G.; Ortiga-Carvalho, T.M.; et al. Gestational age-dependent gene expression profiling of ATP-binding cassette transporters in the healthy human placenta. J. Cell. Mol. Med. 2018, 610–618, doi:10.1111/jcmm.13966.

18. Liong, S.; Lappas, M. Lipopolysaccharide and double stranded viral RNA mediate insulin resistance and increase system a amino acid transport in human trophoblast cells in vitro. Placenta 2017, 51, 18–27, doi:10.1016/j.placenta.2017.01.124.

19. Jones, H.N.; Jansson, T.; Powell, T.L. IL-6 stimulates system A amino acid transporter activity in trophoblast cells through STAT3 and increased expression of SNAT2. Am. J. Physiol. - Cell Physiol. 2009, doi:10.1152/ajpcell.00195.2009.

20. Aye, I.L.M.H.; Jansson, T.; Powell, T.L. Interleukin-1β inhibits insulin signaling and prevents insulin-stimulated system A amino acid transport in primary human trophoblasts. Mol. Cell. Endocrinol. 2013, doi:10.1016/j.mce.2013.07.013.

21. Connor, K.L.; Kibschull, M.; Matysiak-Zablocki, E.; Nguyen, T.T.-T.N.; Matthews, S.G.; Lye, S.J.; Bloise, E. Maternal malnutrition impacts placental morphology and transporter expression: An origin for poor offspring growth. J. Nutr. Biochem. 2020, 108329, doi:10.1016/j.jnutbio.2019.108329.

22. Festing, M.F.W. Design and statistical methods in studies using animal models of development. ILAR J. 2006, 47, 5–14, doi:10.1093/ilar.47.1.5.

23. Coan, P.M.; Angiolini, E.; Sandovici, I.; Burton, G.J.; Constância, M.; Fowden, A.L. Adaptations in placental nutrient transfer capacity to meet fetal growth demands depend on placental size in mice. J. Physiol. 2008, 586, 4567–4576, doi:10.1113/jphysiol.2008.156133.

24. Bloise, E.; Lin, W.; Liu, X.; Simbulan, R.; Kolahi, K.S.; Petraglia, F.; Maltepe, E.; Donjacour, A.; Rinaudo, P. Impaired placental nutrient transport in mice generated by in vitro fertilization. Endocrinology 2012, 153, 3457–3467, doi:10.1210/en.2011-1921.

25. Bloise, E.; Bhuiyan, M.; Audette, M.C.; Petropoulos, S.; Javam, M.; Gibb, W.; Matthews, S.G. Prenatal Endotoxemia and Placental Drug Transport in The Mouse: Placental Size-Specific Effects. PLoS One 2013, 8, doi:10.1371/journal.pone.0065728.

26. Livak KJ1, S.T. Analysis of relative gene expression data using real-time quantitative PCR and the 2(-Delta Delta C(T)) Method. Methods. 2001, 25(4), 402–8.

27. Dyer, B.& Canadian Journal of Biochemistry and Physiology. Can. J. … 1963, 37.

28. Ruiz, J.I.; Ochoa, B. Quantification in the subnanomolar range of phospholipids and neutral lipids by monodimensional thin-layer chromatography and image analysis. J. Lipid Res. 1997, 38, 1482–1489.

29. Murray, S.A.; Morgan, J.L.; Kane, C.; Sharma, Y.; Heffner, C.S.; Lake, J.; Donahue, L.R. Mouse gestation length is genetically determined. PLoS One 2010, doi:10.1371/journal.pone.0012418.

30. Hayward, C.E.; Lean, S.; Sibley, C.P.; Jones, R.L.; Wareing, M.; Greenwood, S.L.; Dilworth, M.R. Placental adaptation: What can we learn from Birthweight:placental weight ratio? Front. Physiol. 2016, 7, 1–13, doi:10.3389/fphys.2016.00028.

31. Khan, R.N.; Hay, D.P. A clear and present danger: Inflammasomes DAMPing down disorders of pregnancy. Hum. Reprod. Update 2015, 21, 388–405, doi:10.1093/humupd/dmu059.

32. Martinelli, L.M.; Reginatto, M.W.; Fontes, K.N.; Andrade, C.B.V.; Monteiro, V.R.S.; Gomes, H.R.; Almeida, F.R.C.L.; Bloise, F.F.; Matthews, S.G.; Ortiga-Carvalho, T.M.; et al. Breast cancer resistance protein (Bcrp/Abcg2) is selectively modulated by lipopolysaccharide (LPS) in the mouse yolk sac. Reprod. Toxicol. 2020, doi:10.1016/j.reprotox.2020.09.001.

33. Martinelli, L.M.; Fontes, K.N.; Reginatto, M.W.; Andrade, C.B.V.; Monteiro, V.R.S.; Gomes, H.R.; Silva-Filho, J.L.; Pinheiro, A.A.S.; Vago, A.R.; Almeida, F.R.C.L.; et al. Malaria in pregnancy regulates P-glycoprotein (P-gp/Abcb1a) and ABCA1 efflux transporters in the Mouse Visceral Yolk Sac. J. Cell. Mol. Med. 2020, 24, 10636–10647, doi:10.1111/jcmm.15682.

34. Toyama RP1, Xikota JC, Schwarzbold ML, Frode TS, Buss Zda S, Nunes JC, Funchal GD, Nunes FC, Walz R, P.M. J Matern Fetal Neonatal Med. 2015, (4), 426–30, doi:10.3109/14767058.2014.918600.

35. Bloise, E.; Feuer, S.K.; Rinaudo, P.F. Comparative intrauterine development and placental function of ART concepti: Implications for human reproductive medicine and animal breeding. Hum. Reprod. Update 2014, 20, 822–839, doi:10.1093/humupd/dmu032.

36. Kim, Y.M.; Romero, R.; Chaiworapongsa, T.; Kim, G.J.; Kim, M.R.; Kuivaniemi, H.; Tromp, G.; Espinoza, J.; Bujold, E.; Abrahams, V.M.; et al. Toll-like receptor-2 and - 4 in the chorioamniotic membranes in spontaneous labor at term and in preterm parturition that are associated with chorioamnionitis. In Proceedings of the American Journal of Obstetrics and Gynecology; 2004; Vol. 191, pp. 1346–1355.

37. Vrachnis, N.; Karavolos, S.; Iliodromiti, Z.; Sifakis, S.; Siristatidis, C.; Mastorakos, G.; Creatsas, G. Impact of mediators present in amniotic fluid on preterm labour. In Vivo (Brooklyn). 2012, 26, 799–812.

38. Shynlova, O.; Tsui, P.; Jaffer, S.; Lye, S.J. Integration of endocrine and mechanical signals in the regulation of myometrial functions during pregnancy and labour. Eur. J. Obstet. Gynecol. Reprod. Biol. 2009, 144, S2, doi:10.1016/j.ejogrb.2009.02.044.

39. Gibb, W. The role of prostaglandins in human parturition. Ann Med. 1998, 30, 235–41, doi:10.3109/07853899809005850.

40. Whittle, W.L.; Holloway, A.C.; Lye, S.J.; Gibb, W.; Challis, J.R.G. Prostaglandin production at the onset of ovine parturition is regulated by both estrogen-independent and estrogen-dependent pathways. Endocrinology 2000, 141, 3783–3791, doi:10.1210/endo.141.10.7703.

41. Chen, Z.; Liu, Q.; Zhu, Z.; Xiang, F.; Wu, R.; Kang, X. Toll-like receptor 4 contributes to uterine activation by upregulating pro-inflammatory cytokine and CAP expression via the NF-κB/P38MAPK signaling pathway during pregnancy. J. Cell. Physiol. 2020, 235, 513–525, doi:10.1002/jcp.28991.

42. Lye, P.; Bloise, E.; Dunk, C.; Javam, M.; Gibb, W.; Lye, S.J.; Matthews, S.G. Effect of oxygen on multidrug resistance in the first trimester human placenta. Placenta 2013, 34, 817–823, doi:10.1016/j.placenta.2013.05.010.

43. Kalabis, G.M.; Kostaki, A.; Andrews, M.H.; Petropoulos, S.; Gibb, W.; Matthews, S.G. Multidrug Resistance Phosphoglycoprotein (ABCB1) in the Mouse Placenta: Fetal Protection1. Biol. Reprod. 2005, 73, 591–597, doi:10.1095/biolreprod.105.042242.

44. Bloise, E.; Petropoulos, S.; Iqbal, M.; Kostaki, A.; Ortiga-Carvalho, T.M.; Gibb, W.; Matthews, S.G. Acute Effects of Viral Exposure on P-Glycoprotein Function in the Mouse Fetal Blood-Brain Barrier. Cell. Physiol. Biochem. 2017, 41, 1044–1050, doi:10.1159/000461569.

45. Lye, P.; Bloise, E.; Nadeem, L.; Peng, C.; Gibb, W.; Ortiga-Carvalho, T.M.; Lye, S.J.; Matthews, S.G. Breast Cancer Resistance Protein (BCRP/ABCG2) Inhibits Extra Villous Trophoblast Migration: The Impact of Bacterial and Viral Infection. Cells 2019, 8, 1150, doi:10.3390/cells8101150.

46. Mason, C.W.; Lee, G.T.; Dong, Y.; Zhou, H.; He, L.; Weiner, C.P. Effect of prostaglandin E2 on multidrug resistance transporters in Human Placental Cells. In Proceedings of the Drug Metabolism and Disposition; American Society for Pharmacology and Experimental Therapy, 2014; Vol. 42, pp. 2077–2086.

47. Silva, J.F.; Serakides, R. Intrauterine trophoblast migration: A comparative view of humans and rodents. Cell Adhes. Migr. 2016, 10, 88–110.

48. Liu, Z.; Liu, W.; Huang, Y.; Guo, J.; Zhao, R.; Yang, X. Lipopolysaccharide significantly influences the hepatic triglyceride metabolism in growing pigs. Lipids Health Dis. 2015, 14, doi:10.1186/s12944-015-0064-8.

49. Lewis, R.M.; Desoye, G. Placental Lipid and Fatty Acid Transfer in Maternal Overnutrition. Ann. Nutr. Metab. 2017, 70, 228–231.

50. Daniel, Z.; Swali, A.; Emes, R.; Langley-Evans, S.C. The effect of maternal undernutrition on the rat placental transcriptome: protein restriction up-regulates cholesterol transport. Genes Nutr. 2016, 11, 1–12, doi:10.1186/s12263-016-0541-3.

51. Cetin, I.; Parisi, F.; Berti, C.; Mandó, C.; Desoye, G. Placental fatty acid transport in maternal obesity. J. Dev. Orig. Health Dis. 2012, 3, 409–414.

52. Hirai, T.; Fukui, Y.; Motojima, K. PPARα; Agonists Positively and Negatively Regulate the Expression of Several Nutrient/Drug Transporters in Mouse Small Intestine. Biol. Pharm. Bull. 2007, 30, 2185–2190, doi:10.1248/bpb.30.2185.

53. Merrell, M.D.; Nyagode, B.A.; Clarke, J.D.; Cherrington, N.J.; Morgan, E.T. Selective and cytokine-dependent regulation of hepatic transporters and bile acid homeostasis during infectious colitis in mice. Drug Metab. Dispos. 2014, 42, 596–602, doi:10.1124/dmd.113.055525.

54. Murakami, M.; Harada, M.; Kamimura, D.; Ogura, H.; Okuyama, Y.; Kumai, N.; Okuyama, A.; Singh, R.; Jiang, J.J.; Atsumi, T.; et al. Disease-association analysis of an inflammation-related feedback loop. Cell Rep. 2013, doi:10.1016/j.celrep.2013.01.028.

55. Zammit, N.W.; Tan, B.M.; Walters, S.N.; Liuwantara, D.; Villanueva, J.E.; Malle, E.K.; Grey, S.T. Low-dose rapamycin unmasks the protective potential of targeting intragraft NF-κB for islet transplants. Cell Transplant. 2013, doi:10.3727/096368912X658737.

56. Coughlin, B.; Schnabolk, G.; Joseph, K.; Raikwar, H.; Kunchithapautham, K.; Johnson, K.; Moore, K.; Wang, Y.; Rohrer, B. Connecting the innate and adaptive immune responses in mouse choroidal neovascularization via the anaphylatoxin C5a and γδT-cells. Sci. Rep. 2016, doi:10.1038/srep23794.

57. Gong, Z.K.; Wang, S.J.; Huang, Y.Q.; Zhao, R.Q.; Zhu, Q.F.; Lin, W.Z. Identification and validation of suitable reference genes for RT-qPCR analysis in mouse testis development. Mol. Genet. Genomics 2014, doi:10.1007/s00438-014-0877-6.

