## Supplementary Figure for "Effect of sublethal prenatal endotoxaemia on murine placental transport systems and lipid homeostasis"

**Supplementary table 1: List of Primers Used in the Present Study.**

| Primer | Sequence | Reference | Genbank no |
|---|---|---|---|
| ***Abca1*** | 5'GCAGATCAAGCATCCCAACT3'<br>3'CCAGAGAATGTTTCATTGTCCA5' | [52] | NM_013454.3 |
| ***Abcb1a*** | 5'GGGCATTTACTTCAAACTTGTCA3'<br>3'TTTACAAGCTTCATTTCCTAATTCAA5' | [52] | NM_011076.3 |
| ***Abcb1b*** | 5'AAGCCAGTATTCTGCCAAGCAT3'<br>3'CTCCAGACTGCTGTTGCTGATG5' | [52] | NM_011075.2 |
| ***Abcb4*** | 5'GAAGGGATCTACTTCAGACTCGTT3'<br>3'TCAACTTCAAATTCTTCTGACAGG5' | [52] | NM_008830.2 |
| ***Abcc2*** | 5'TAATGAGGCGCCGTGGGTGAC3'<br>3'GTCCTGCCCACCACACCGAC5' | [52] | NM_013806.2 |
| ***Abcc5*** | 5'AAATGTATGCCTGGGTCAAAGC3'<br>3'TGGCGATCACTACCACAATAGG5' | * | NM_013790.2 |
| ***Abcf2*** | 5'TGTCCACATTATCAACCTCTCCC 3'<br>3'TCACGTTTCCCAATAGCCGAG 5' | * | NM_013853.2 |
| ***Abcg1*** | 5'GCTCCATCGTCTGTACCATCC3'<br>3'ACGCATTGTCCTTGACTTAGG5' | * | |
| ***Abcg2*** | 5'GCCTTGGAGTACTTTGCATCA3'<br>3'AAATCCGCAGGGTTGTTGTA5' | [53] | NM_011920.3 |
| ***Il6*** | 5'GAGGATACCACTCCCAACAGACC 3'<br>3'AAGTGCATCATCGTTGTTCATACA 5' | [54] | NM_031168.2 |
| ***Cxcl1*** | 5'ACCCGCTCGCTTCTCTGT 3'<br>3'AAGGGAGCTTCAGGGTCAAG 5' | [54] | NM_008176.3 |
| ***Ccl2*** | 5'GGTCCCTGTCATGCTTCTGG 3'<br>3'CCTGCTGCTGGTGATCCTCT 5' | [55] | NM_011333.3 |
| ***Pparg*** | * | * | NM_001127330.2 |
| ***CD36(FAT)*** | 5'CGCAGCCTCCTTTCCACCTTTTGT3'<br>3' TGGTTGTCTGGATTCTGGAGGGGT5' | * | NM_001159556.1 |

| Primer | Sequence | Reference | Genbank no |
|---|---|---|---|
| ***Fatp1*** | 5'CGCTTTCTGCGTATCGTCTG3'<br>3'GATGCACGGGATCGTGTCT5' | * | NM_011977.4 |
| ***Fabppm*** | 5'GCGTCCCGAGCAGTGGAAGGA3'<br>3'GGCATACGATTGGCAGAGGCAGA5' | * | NM_010325.2 |
| ***Lpl*** | 5'TGGCGTAGCAGGAAGTCTGA3'<br>5'TGCCTCCATTGGGATAAATGTC3' | * | NM_008509.2 |
| ***B2m*** | 5'TTCTGGTGCTTGTCTCCAGTATGTTC3'<br>5'GCTTCCCATTCACTGA3' | * | NM_011149.2 |
| **B-actina (Actb)** | 5'AAATCTGGCACCACACCTTC3'<br>5'GGGGTGTTGAAGGTCTCAAA3' | [56] | NM_007393.5 |
| ***Gapdh*** | 5'TGTGTCCGTCGTGGATCTGA 3'<br>3'TTGCTGTTGAAGTCGCAGGAG 5' | [57] | NM_001289726.1 |
| ***Ywhaz*** | 5' GAAAAGTTCTTGATCCCCAATGC 3'<br>5' TGTGACTGGTCCACAATTCCTT 3' | * | NM_011740.3 |

*Gene specific primers were designed with primer-BLAST (http://www.ncbi.nlm.gov/tools/primer-blast).

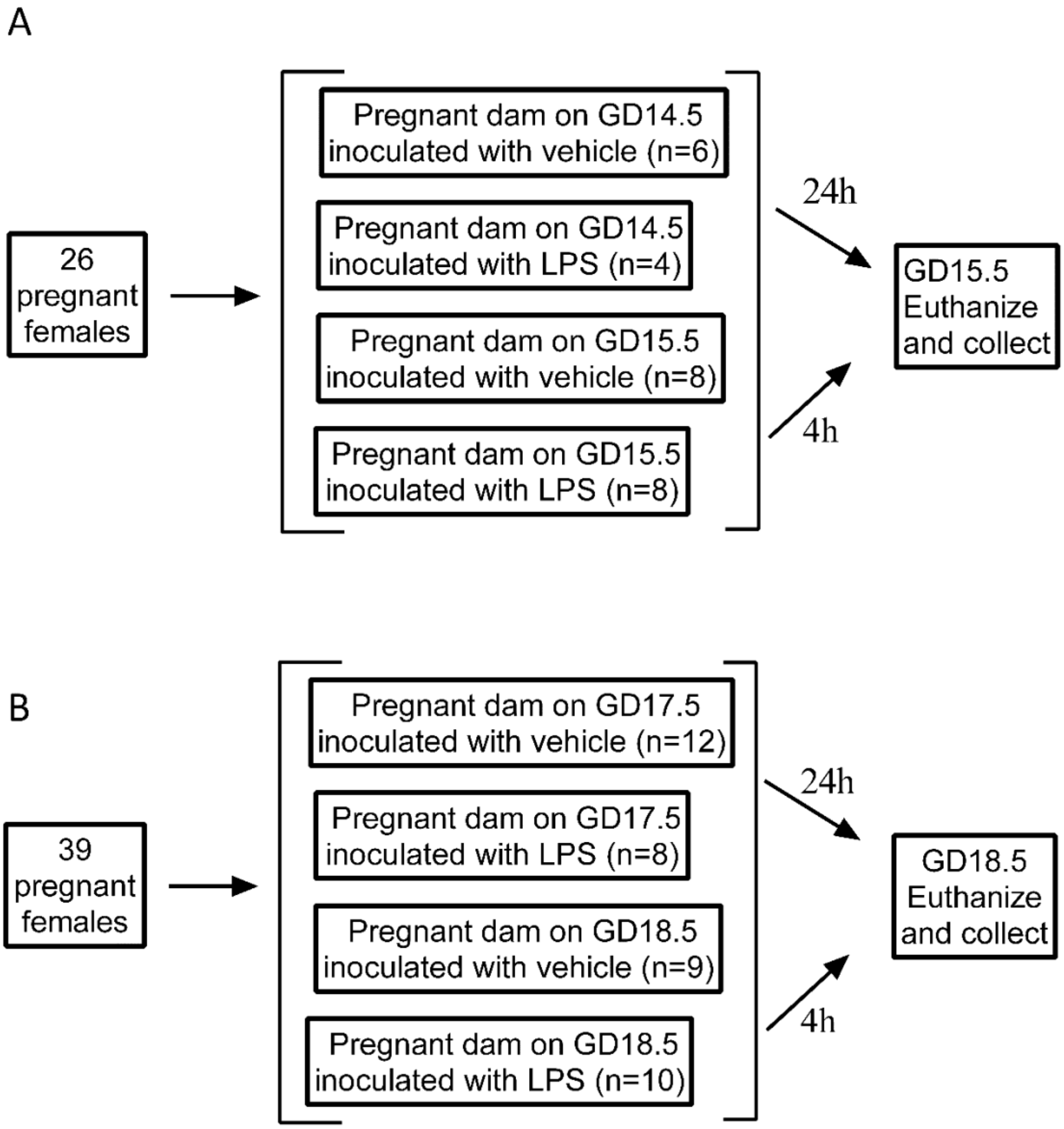


**Supplementary figure 1: Study Design.**

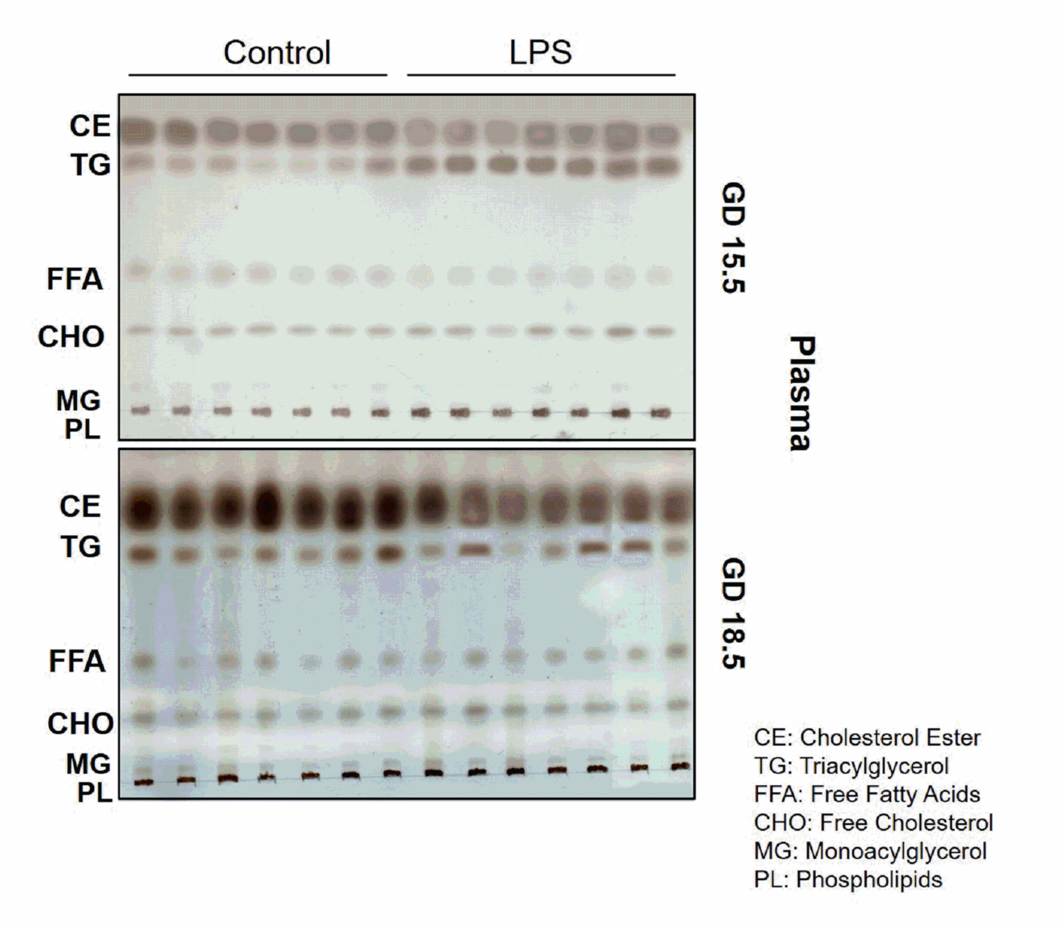


**Supplementary figure 2: Representative image of TLC plate from maternal plasma at GD 15.5 and 18.5.** 15.5/4h: n=6 (control group); n=5 (LPS group). CE: Cholesterol Ester; TG: Triacylglycerol; FFA: Free Fatty Acids; CHO: Free Cholesterol; MG: Monoacylglycerol; PL: Phospholipids.

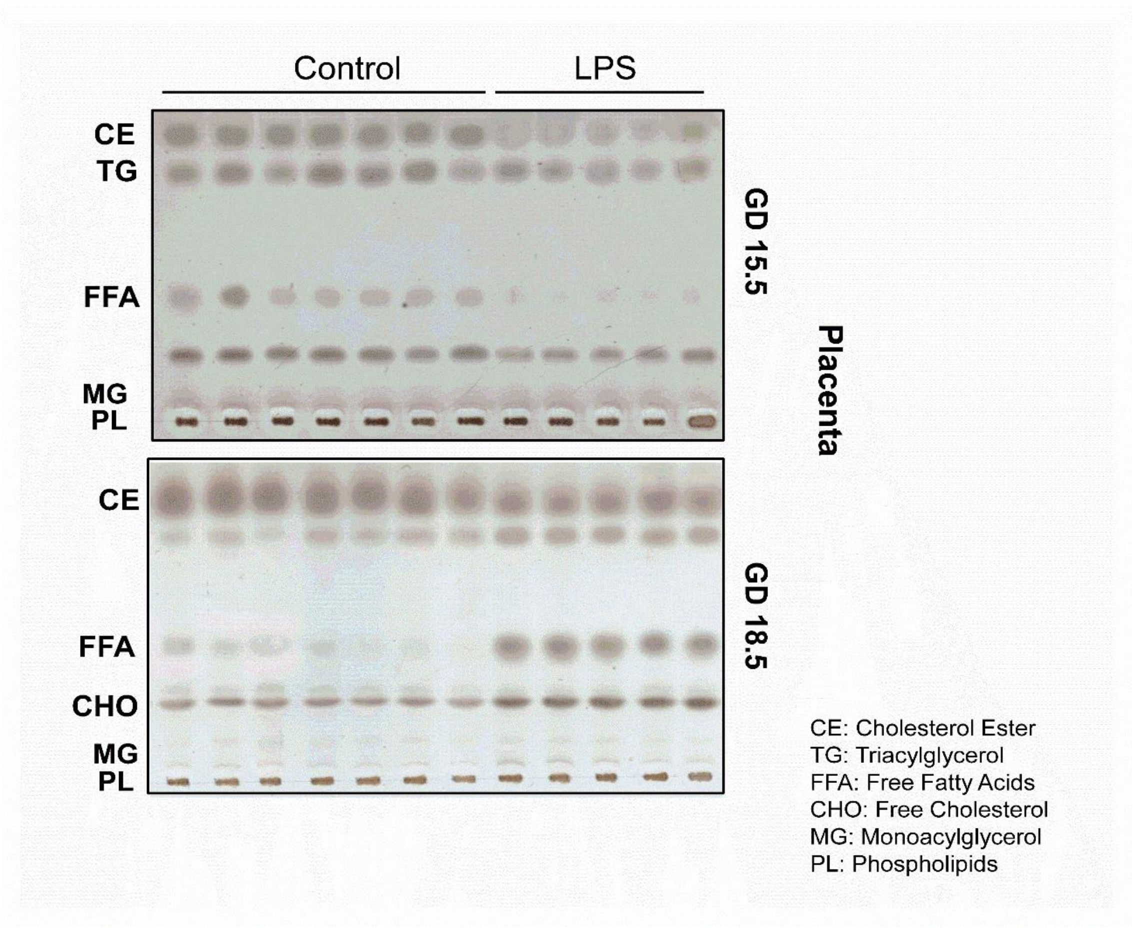


**Supplementary figure 3: Representative image of TLC plate from placenta at GD 15.5 and 18.5.** 15.5/4h: n=6 (control group); n=5 (LPS group); 18.5/4h: n=9 (control group); n=7 (LPS group). CE: Cholesterol Ester; TG: Triacylglycerol; FFA: Free Fatty Acids; CHO: Free Cholesterol; MG: Monoacylglycerol; PL: Phospholipids.

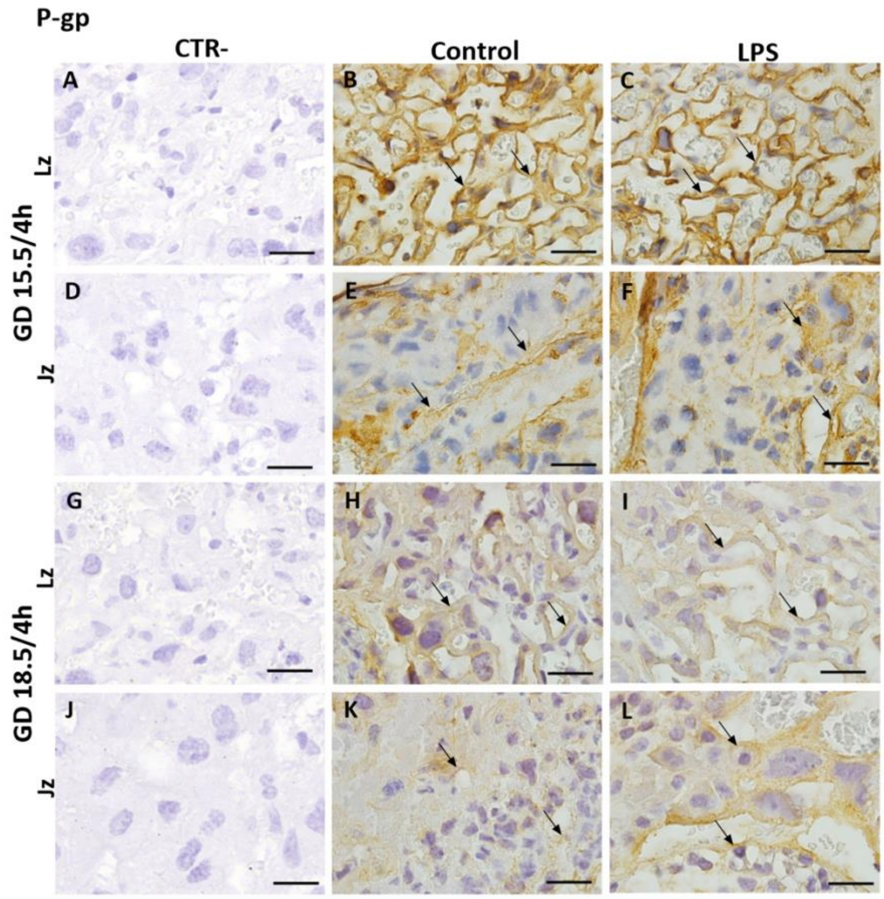


**Supplementary figure 4: Representative photomicrographies of immunohistochemistry staining of P-gp in higher magnification 4 h after LPS insult at GD15.5 and 18.5 in the labyrinth (Lz) and junctional (Jz) zones of the placenta.** n= 5/group. Scale bar =50 μm. Normal serum was incubated with negative control sections instead of the P-gp primary antibody.

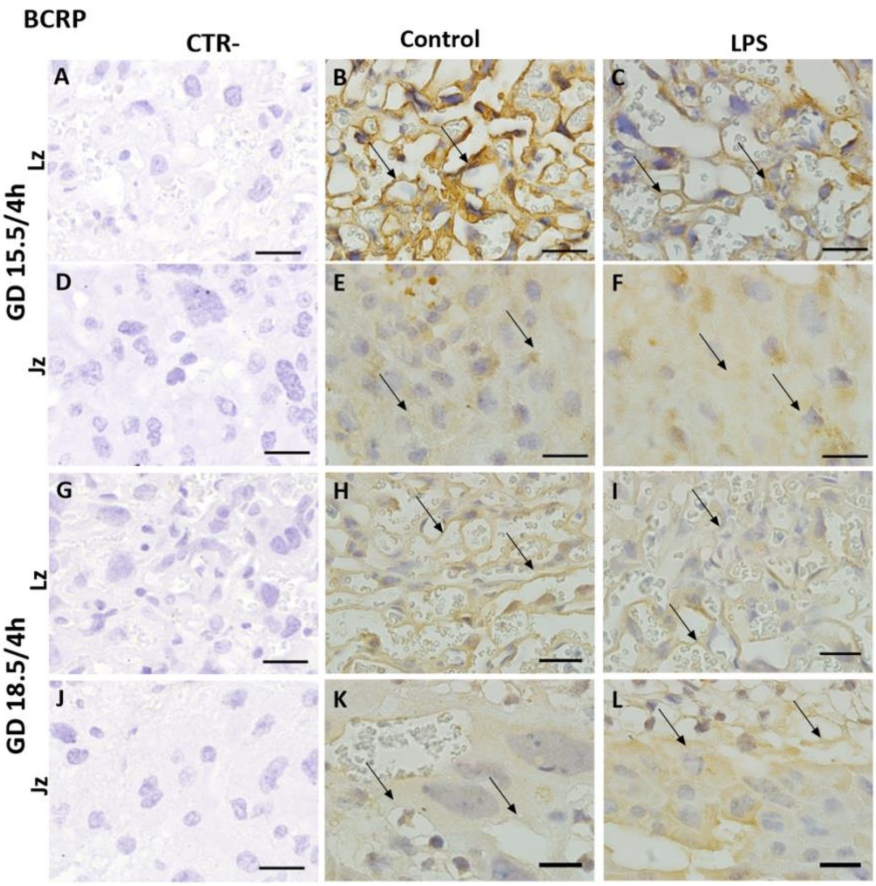


**Supplementary figure 5: Representative photomicrographies of immunohistochemistry staining of Bcrp in higher magnification 4 h after LPS insult at GD15.5 and 18.5 in the labyrinth (Lz) and junctional (Jz) zones of the placenta.** n= 5/group. Scale bar =50 µm. Normal serum was incubated with negative control sections instead of the Bcrp primary antibody.

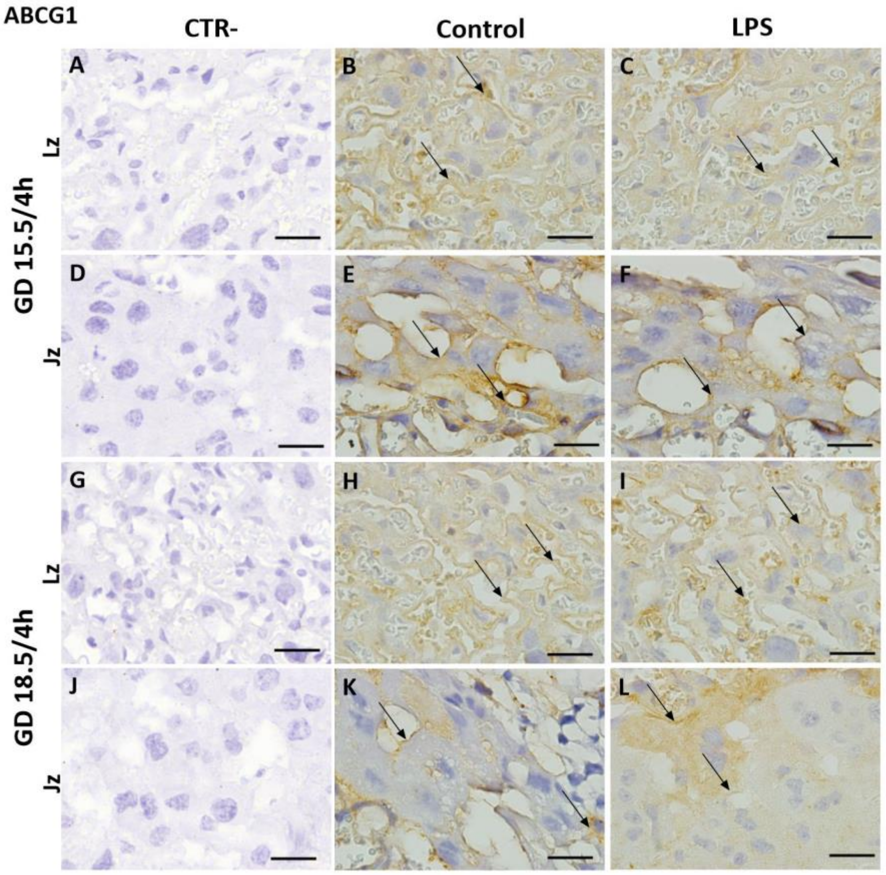


**Supplementary figure 6: Representative photomicrographies of immunohistochemistry staining of Abcg1 in higher magnification 4 h after LPS insult at GD15.5 and 18.5 in the labyrinth (Lz) and junctional (Jz) zones of the placenta.** n= 5/group. Scale bar =50 µm. Normal serum was incubated with negative control sections instead of the Abcb1 primary antibody.
